# Aspartate transaminases are required for blood development

**DOI:** 10.1101/2025.09.16.675844

**Authors:** Narges Pourmandi, Greggory Myers, Arjun Jha, Kelsey Temprine, Amanda Sankar, Nupur K. Das, Ridwana Khan, Siva Kumar Natarajan, Cristina Castillo, Peter Sajjakulnukit, Noah S. Nelson, Matthew D. Perricone, Indrani Talukder, Aaron D. denDekker, Lin Lin, Dominik Awad, Wesley Huang, Lei Yu, Navdeep S. Chandel, Rami Khoriaty, Yatrik M. Shah, Costas A. Lyssiotis

## Abstract

Red blood cells (RBCs) have a limited lifespan of approximately 120 days. This necessitates continuous RBC production, resulting in ∼200 billion new RBCs made per day to maintain oxygen delivery. Despite this enormous biosynthetic demand, the metabolic pathways supporting erythropoiesis are poorly understood. We profiled metabolites across four independent models of elevated erythropoiesis and a consistent increase in aspartate levels emerged when compared to controls. This suggested a potential role for aspartate metabolism in RBC production. To test this, we deleted the aspartate aminotransferases *Got1* or *Got2* globally or in an erythroid-specific manner. Loss of either enzyme resulted in anemia, with *Got2* deficiency producing a more severe phenotype. Individual loss of either *Got* or dual *Got* deletion led to an erythroid defect, where early progenitors accumulated. In human and mouse models of erythropoiesis, GOT1 and GOT2 loss had opposing impacts on aspartate despite exhibiting similar anemic phenotypes, suggesting an aspartate-independent function for these enzymes. GOT1 and GOT2 are also components of the malate-aspartate shuttle (MAS), which regulates NAD(H) homeostasis. However, conditional deletion of another MAS enzyme, *Mdh1*, did not cause anemia, and alleviating NADH reductive stress in GOT2-deficient cells with cytoplasmic bacterial NADH oxidase (LbNOX) failed to restore erythropoiesis. Instead, transcriptomic and epigenetic analyses revealed dysregulation of chromatin histone modifications in GOT-deficient erythroid cells, implicating epigenetic dysfunction as a driver of defective erythropoiesis. Collectively, these findings identify a previously unrecognized role for GOT1 and GOT2 as a central link between metabolism and epigenetic regulation during erythroid development. These insights may inform the development of new therapeutic strategies for anemia.

## Introduction

Red blood cells (RBCs) are the most abundant cell type in the human body.^1^ To sustain life, humans require oxygen, and RBCs have evolved as a highly specialized cell type optimized for oxygen transport. The production of RBCs through erythropoiesis is a multistep process that can be divided into 2 stages (**Fig. 1a**).^2^ The first phase, early erythropoiesis, encompasses the commitment of multipotent hematopoietic stem cells in the bone marrow to the erythroid lineage and subsequent production of erythroid restricted progenitors called burst-forming units-erythroid (BFU-E) and colony-forming units-erythroid (CFU-E) cells. In the second phase, terminal erythroid differentiation, proerythroblasts (ProEbs) undergo major morphological and functional changes that are unique to their terminal function of oxygen transport. This includes the loss of their nuclei and mitochondria and massive production of hemoglobin. Ultimately, following several cell divisions and intermediary forms, ProEbs become reticulocytes that are expelled from the bone marrow to the circulating blood, where they mature to form RBCs.

**Fig. 1:**
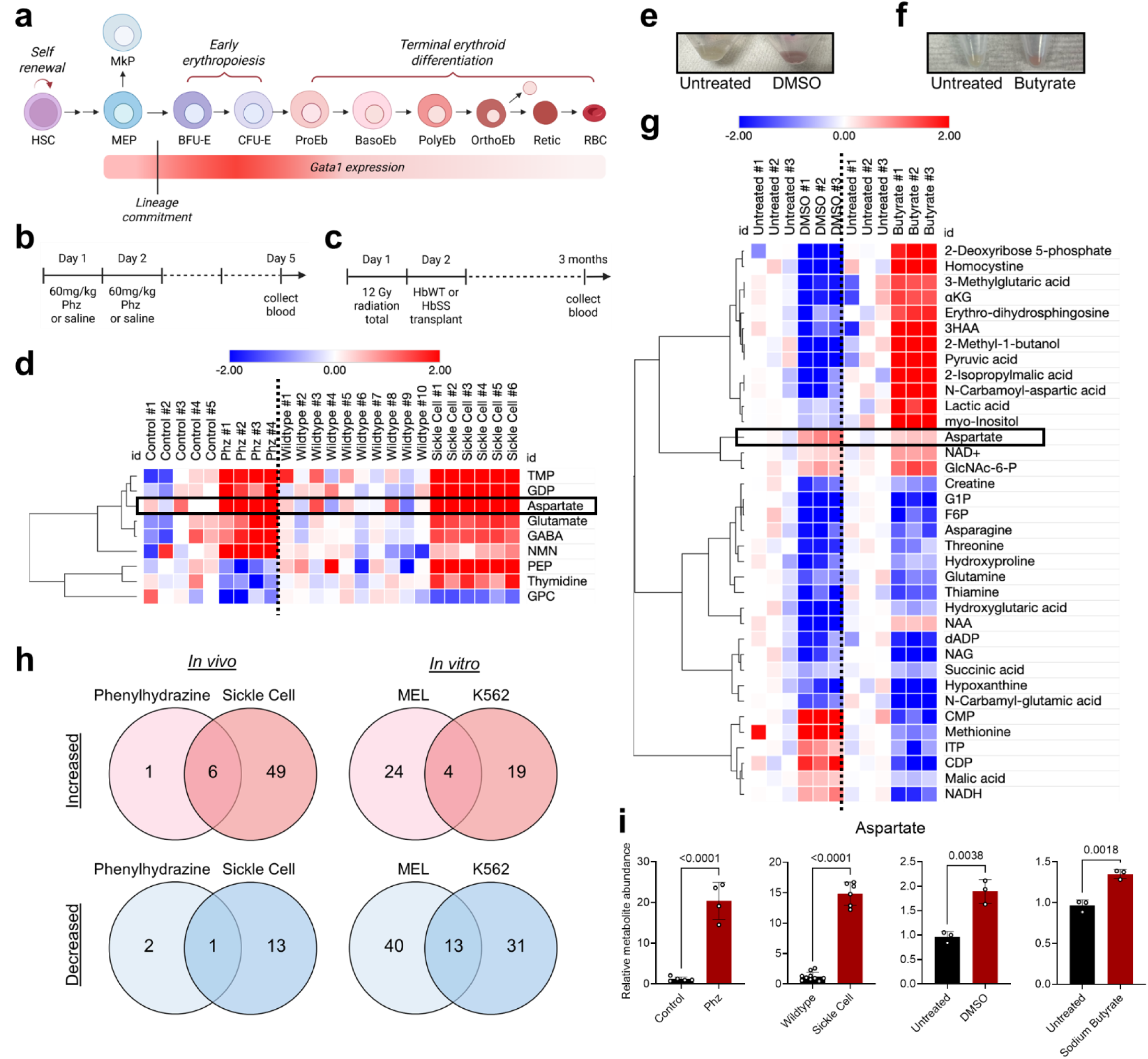
Four diverse models of erythropoiesis exhibit elevated aspartate. a, Stages of erythroid differentiation starting from hematopoietic stem cells (HSC), which, following several cell divisions and intermediary progenitor cells, generate megakaryocyte-erythroid progenitors (MEPs). MEPs give rise to either a megakaryocyte progenitor (MkP) cell or a burst-forming unit–erythroid (BFU-E) cell. BFU-Es sequentially generate colony-forming unit–erythroid (CFU-E) cells, proerythroblasts (ProEb), basophilic erythroblasts (BasoEb), polychromatic erythroblasts (PolyEb), orthochromatic erythroblasts (OrthoEb), anucleate reticulocytes (Retic), and finally mature red blood cells (RBCs). **b,** Experimental timeline of control mice treated with saline (n=6) and mice treated with phenylhydrazine (Phz, n=6). **c,** Experimental timeline of wildtype (WT) mice that received either WT bone marrow (n=9) or sickle cell disease bone marrow (n=7). **d,** Heat map of the overlapping significantly changed metabolites (p-value <0.05) in the circulating RBCs from phenylhydrazine (Phz)-treated mice and sickle cell disease mice. The log_2_ of the median centered to control metabolite abundances are shown. **e,** MEL differentiation with 2% DMSO for 4 days shows red pellet, reflective of increased hemoglobin production. **f,** K562 differentiation with 1.5mM sodium butyrate for 4 days shows red pellet, reflective of increased hemoglobin production. **g,** Heat map of the overlapping significantly changed metabolites (p-value <0.05) in the DMSO treated MEL cells and sodium butyrate treated K562 cells. The log_2_ of the median centered to control metabolite abundances are shown. **h,** Overlap in the significantly increased (top row) or decreased (bottom row) metabolites (p-value <0.05) between the Phz-treated mice and sickle cell disease mice (left) and between the DMSO treated MEL cells and sodium butyrate treated K562 cells (right). **i,** Relative abundances of aspartate (median centered to the control conditions) in the aforementioned models. The statistical significance test for all graphs was done with an unpaired two-tailed t-test with a significance threshold level of 0.05.

Anemia is a condition of reduced healthy RBCs, resulting in diminished oxygen delivery to tissues and manifesting as fatigue, shortness of breath, and/or tachycardia^3^. Approximately 1 in 4 people have anemia globally, with females and children under five years of age disproportionally affected. Anemia can result from defects in proliferation of erythroid precursors, inefficient erythroid maturation, or loss of RBCs from circulation.^4^ Despite differences in pathophysiology among these causes of anemia, the mainstay therapies for most anemias address symptomatic control with RBC transfusions. Transfusions lead to an accumulation of excess iron that, in turn, can result in tissue iron overload and clinical consequences, such as liver cirrhosis, diabetes mellitus, and heart failure.^5,6^ Recently, the TGFβ inhibitor luspatercept was found to benefit patients with certain dyserythropoietic conditions such as myelodysplastic syndromes and beta-thalassemia; however, the benefit is often modest and relatively short lived.^7,8^ Thus, there is a clear unmet need to develop targeted therapies that address the underlying causes of anemia.^4,9^

Erythropoiesis is a metabolically intensive process, with erythroid progenitors generating over 2 million new RBCs every second.^10^ Yet, only in the past decade has metabolism emerged as a possible therapeutic avenue. This is exemplified in sickle cell disease, in which erythrocytes have increased susceptibility to oxidant damage and thus rely on the NADPH pool for antioxidant precursors.^11^ The amino acid glutamine, a precursor in the regulation of NADPH, was found to be decreased in patients with sickle cell disease.^11^ As a result, glutamine was evaluated in a clinical trial (NCT01179217) as a therapy for sickle cell disease, and it modestly reduced the number of sickle cell crises compared to placebo.^12^ More recently, mitapavit, a pyruvate kinase activator, is being explored for treatment of sickle cell anemia.^13^ Although the clinical benefit of these metabolic therapies have been modest, they highlight the potential of targeting metabolism to develop better therapies for erythroid disorders.

Over the past decade, significant progress has been made in understanding aspects of metabolism that regulate terminal erythroid differentiation. We now understand that hemoglobin production is promoted by amino acid-induced mTOR signaling^14^. As a part of heme production, exogenous glutamine is converted to α-ketoglutarate (aKG), which is then used to produce succinyl-CoA.^15^ Moreover, late-stage erythropoiesis is dependent on decreasing αKG-driven oxidative phosphorylation^16^. Furthermore, glutamate conversion to glutamine is essential during terminal erythropoiesis for ammonia detoxification, a highly abundant byproduct of heme synthesis.^17^

In contrast, the metabolic pathways that regulate HSC commitment to the erythroid lineage remain less well understood. Thus far, it has been established that glutamine-dependent de novo nucleotide biosynthesis is required for erythroid commitment under conditions of stress, such as hemolytic anemia.^18^ Moreover, it has been shown that erythroid commitment requires increased oxidative phosphorylation to sustain erythroid differentiation, and this is driven by anaplerotic utilization of glutamine to produce αKG.^16^

The malate-aspartate shuttle (MAS) is an evolutionarily conserved metabolic pathway that maintains redox and energy homeostasis between the cytosol and the mitochondria. Energy generated in the cytosol from glycolysis as “NADH” cannot be directly transferred into the mitochondria. Instead, the MAS transfers this energy stored within metabolites (i.e. malate and aspartate), which are able to cross the mitochondrial membrane. The MAS is composed of two pyridoxal 5’-phosphate (PLP) dependent aspartate transaminases (cytosolic GOT1 and mitochondrial GOT2) and two malate dehydrogenases (cytosolic MDH1 and mitochondrial MDH2). Independent of their functions in the MAS, the GOT enzymes are the only cellular source of aspartate, a key biosynthetic intermediate in nucleotide and amino acid biosynthesis.^19–22^ In the process of regulating aspartate levels, the GOTs also control and are an important source of αKG regulation. Thus, the GOT enzymes can serve multiple simultaneous functions, in bioenergetics and redox balance (coupling glycolysis to respiration), biosynthesis (making aspartate and αKG), and regulating gene expression (through αKG).

In studying the metabolic determinants of erythropoiesis, we observed that four independent models of increased erythropoiesis were met with increased aspartate levels. Thus, to determine the role of the GOTs in erythropoiesis, we crossed our tamoxifen-inducible *Gata1*-CreERT2 allele^23^ with our conditional *Got* alleles to knockout the GOTs during erythropoiesis. Erythroid-specific *Got1* or *Got2* deletion resulted in anemia, with an erythroid differentiation defect observed at an early erythroid progenitor stage. These results are in marked contrast to findings in hematopoietic stem cells (HSCs), where opposite effects are observed for *Got1* loss and *Got2* loss, resulting in enhanced and reduced HSC function, respectively.^24^ The disparate effects of GOT1 or GOT2 deficiency on HSC function are ascribed to opposite aspartate biosynthetic activities for the GOT enzymes, rather than their roles in αKG production or energy transfer within the MAS. We hypothesized that the similar effects of *Got* deletion in erythropoiesis may be attributed to their central role in the MAS and redox balance. To test this, we generated mice to delete malate dehydrogenase 1 (MDH1), another enzyme in the MAS. This deletion would also lead to a dysfunctional MAS and NADH reductive stress, but this resulted in adequate erythropoiesis with no anemia. To validate this further, we crossed a conditional cytoplasmic *Lactobacillus brevis* NADH oxidase (*Lb*NOX) to the *Gata1*-CreERT2 mice. *Lb*NOX is a bacterial enzyme that can ameliorate NADH reductive stress, but despite its expression, the anemic phenotype persisted. This then led us to hypothesize that the similar roles of the GOT enzymes may be ascribed to their influence on αKG balance, which is known to impact epigenetic regulation.^25^ Indeed, analysis of chromatin histone markers led us to find decreased methylation of several activating histone modifications, suggestive of enhanced gene repression particularly in *GOT2* deleted cells. Our findings uncover a novel role for the GOTs during early erythropoiesis in a mechanism that is independent from the MAS and aspartate biosynthesis and instead may be dependent on epigenetic regulation.

## Results

### Four diverse models of erythropoiesis exhibit elevated aspartate

To begin defining the metabolic changes that occur during erythropoiesis (**Fig. 1a**), we employed targeted liquid chromatography-coupled mass spectrometry (LC/MS)-based metabolomic analysis^22,26^ in two *in vivo* mouse models of increased erythropoiesis and two *in vitro* models of erythroid differentiation. First, we induced hemolytic anemia in adult wildtype (WT) mice by phenylhydrazine (Phz) administration (**Fig. 1b**). Phz produces reactive oxygen species in RBCs that lead to lipid peroxidation and cell lysis.^27,28^ Second, using an orthogonal model of hemolytic anemia, we transplanted total bone marrow cells harvested from sickle cell disease mice into lethally irradiated WT recipient mice (**Fig. 1c**). Blood was collected from the mice in these two models and the RBC count was confirmed to be decreased in both the Phz-treated mice and the transplanted sickle cell disease mice when compared to their controls (**Extended Data Fig. 1a,b**). Next, metabolites were extracted from the circulating RBCs, and metabolomic analysis showed a 16.8-fold and a 12-fold increase in aspartate levels in the RBCs of Phz-treated and transplanted sickle cell disease mice, respectively, when compared to their controls (**Fig. 1d, Extended Data Fig. 2a,b**).

To determine whether the metabolomic changes were cell autonomous, two *in vitro* models of erythroid differentiation were assessed. The first model was the MEL cell line, a transformed murine spleen red cell precursors arrested at the proerythroblast stage^29^ with the potential to undergo heme biosynthesis with addition of DMSO.^29^ The second model was the K562 cell line, which is a human chronic myelogenous leukemia cell line^30^ with the potential to undergo erythroid differentiation with addition of sodium butyrate^31^. Increased hemoglobinization was observed visually in each cell line 4 days post treatment, suggesting successful differentiation into erythroid cells (**Fig. 1e,f**). Quantitative PCR analysis also showed a significant increase in β-globin expression, further supporting erythroid differentiation (**Extended Data Fig. 1c,d**). Metabolomic analysis of the cells showed a 2-fold and 1.4-fold aspartate increase in erythroid differentiated MEL and K562 cells compared to untreated control cells (**Fig. 1g-i, Extended Data Fig. 2c,d**). Given the elevation in aspartate in four distinct models of erythropoiesis, we hypothesized that erythropoiesis necessitates an increase in aspartate synthesis or a decrease in aspartate consumption.

Aspartate is a nonessential amino acid that can be produced by all cells^32^, predominantly via the mitochondrial aspartate aminotransferase, GOT2.^20,21,33,34^ Aspartate exists in low levels in circulation^35^ and its uptake requires a specialized transporter whose expression is largely restricted to neural lineages. This indicates that aspartate is primarily made in cells to meet demand. In addition to its role in protein biosynthesis, aspartate is required for de novo pyrimidine and purine synthesis.^36^ Moreover, aspartate serves an important role in the MAS, which facilitates transfer of glycolytic energy in the form of NADH from the cytosol into the mitochondria for oxidative phosphorylation (**Fig. 2a**).^34^ Since NADH cannot cross the mitochondrial membrane, its electrons are placed on malate, which enters through a transporter.^34^ Within this shuttle, GOT2 uses oxaloacetate and glutamate to generate aspartate and αKG (**Fig. 2a**). Aspartate and αKG are then transported into the cytoplasm to be utilized by the cytosolic isoform GOT1 to generate glutamate and oxaloacetate. In instances where there is demand for GOT2-mediated aspartate or αKG biosynthesis, such as in proliferating progenitors, these GOT2 products are used to support other cellular biosynthetic pathways outside of the MAS.^37^

**Fig. 2:**
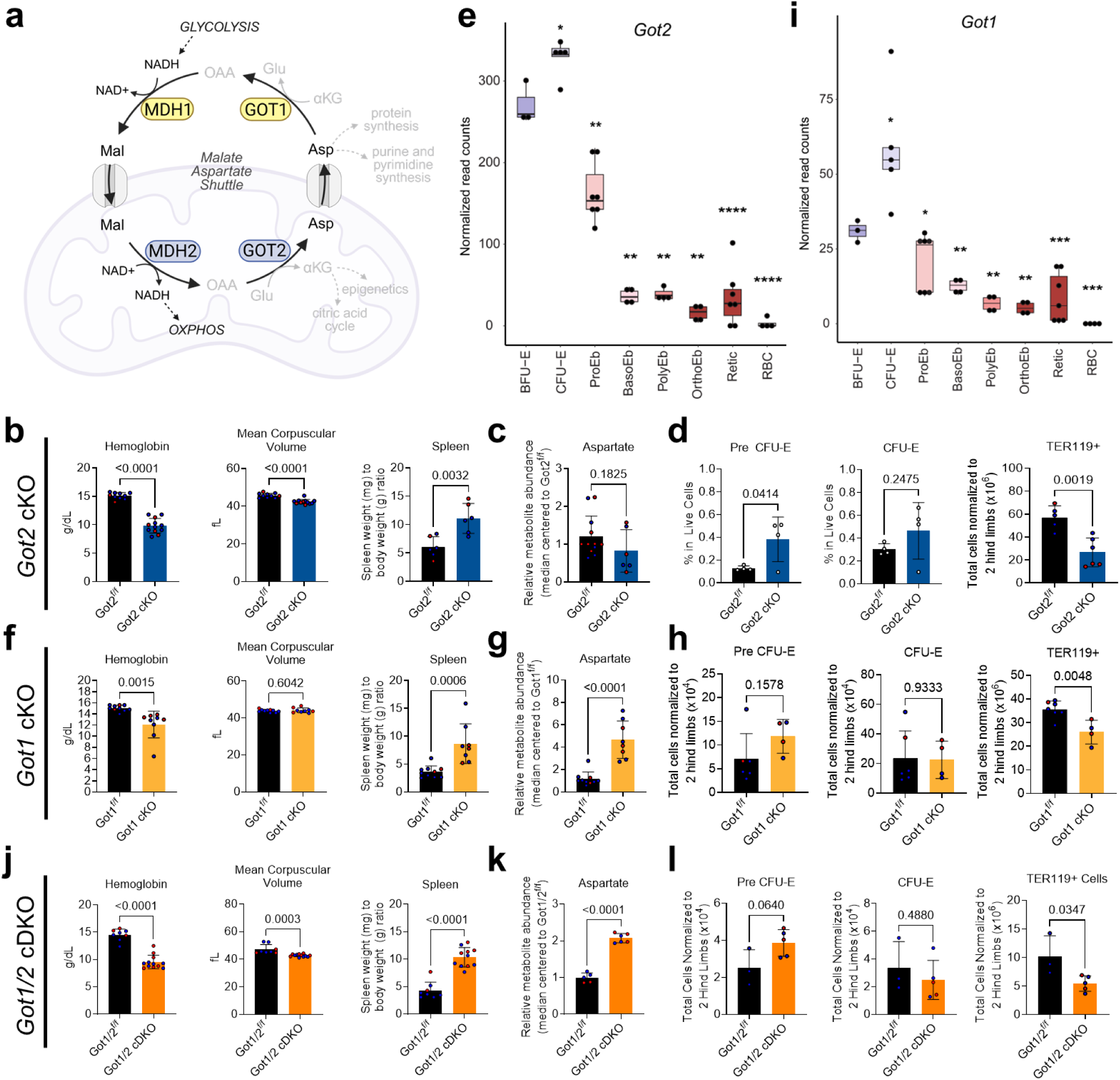
Erythroid-specific deletion of *Got2* and *Got1* in adult mice results in anemia and elevated erythroid-restricted progenitors. a, The malate-aspartate shuttle moves reducing equivalents from NADH made during glycolysis onto malate for transport into the mitochondria for oxidative phosphorylation (OXPHOS). *Slc25a11* encodes the αKG-malate antiporter that moves malate into the mitochondria, and *Slc25a12* and *Slc25a13* encode the glutamate-aspartate antiporters that move aspartate out of the mitochondria. Mal: malate; Asp: aspartate; Glu: glutamate; OAA: oxaloacetate; αKG: α-ketoglutarate. **b,** Hemoglobin and mean corpuscular volume of *Got2^f/f^* control and *Gata1-CreERT2;Got2^Δ/Δ^*(*Got2* cKO) mice after 1 month of tamoxifen, with data presented from 2 independent experiments. The spleen weight normalized to body weight after 1 month of tamoxifen. **c,** Relative abundance of aspartate in circulating RBCs of control and *Got2* cKO mice. Data presented are from 3 independent experiments. **d,** Frequency of pre CFU-E and CFU-E cells (early erythropoiesis) in the bone marrow. Total count of TER119+ cells (terminal erythroblasts) in the bone marrow. **e,** The relative mRNA expression of *Got2* during consecutive cell stages of mouse erythropoiesis, shown as normalized mRNA read counts by RNA-seq. The statistical significance test was done with an unpaired two-tailed t-test comparing the normalized mRNA read counts of the cell stage with those in the BFU-E stage. * p<0.05; ** p<0.01; *** p<0.001; **** p<0.0001. **f,** Hemoglobin and mean corpuscular volume of *Got1^+/f^* and *Got1^f/f^* control and *Gata1-CreERT2;Got1^Δ/Δ^*(*Got1* cKO) mice after 1 month of tamoxifen, with data presented from 2 independent experiments. The spleen weight normalized to body weight after 2 months of tamoxifen. **g,** Relative abundance of aspartate in circulating RBCs of control and *Got1* cKO mice. Data presented are from 2 independent experiments. **h,** Total counts of pre CFU-E and CFU-E cells (early erythropoiesis) in the bone marrow. Total counts of TER119+ cells (terminal erythroblasts) in the bone marrow. **i,** The relative mRNA expression of *Got1* during consecutive cell stages of mouse erythropoiesis, shown as normalized mRNA read counts by RNA-seq. The statistical significance test was done with an unpaired two-tailed t-test comparing the normalized mRNA read counts of the cell stage with those in the BFU-E stage. * p<0.05; ** p<0.01; *** p<0.001; **** p<0.0001. **j,** Hemoglobin and mean corpuscular volume of *Got1/2^f/f^*control and *Gata1-CreERT2;Got1^Δ/Δ^Got2^Δ/Δ^* mice (*Got1/2* cDKO) after 1 month of tamoxifen, with data presented from 2 independent experiments. The spleen size normalized to body weight after 2 months of tamoxifen. **k,** Relative abundance of aspartate in circulating RBCs of control and *Got1/2* cDKO mice. **(L)** Total cell count of pre CFU-E and CFU-E cells (early erythropoiesis) in the bone marrow. Total cell count of TER119+ cells (terminal erythroblasts) in the bone marrow. Red and blue circles represent female and male mice, respectively. The statistical significance test for all graphs was done with an unpaired two-tailed t-test with a significance threshold level of 0.05.

To test the necessity of aspartate biosynthesis in erythropoiesis, we generated whole-body *Got2* knock out (KO) mice by breeding *Got2* floxed mice^22^ with mice expressing a tamoxifen-inducible Cre recombinase from the ubiquitin promoter, *Ub-Cre-ERT2*.^38^ Eight to twelve week old *Ub-CreERT2;Got2^Δ/Δ^* and control mice were placed on tamoxifen for 1 month (**Extended Data Fig. 3a**). *Got2* deletion was confirmed by evaluating cellular protein and messenger RNA levels in the liver (**Extended Data Fig. 3b**). *Ub-CreERT2;Got2^Δ/Δ^* mice had microcytic anemia and spleen weights that trended higher in the *Ub-CreERT2;Got2^Δ/Δ^* mice (**Extended Data Fig. 3c**). Moreover, the anemia observed in *Ub-CreERT2;Got2^Δ/Δ^* mice was not due to a defect in oxygen sensing or EPO production as the circulating EPO level was significantly elevated in *Ub-CreERT2;Got2^Δ/Δ^* mice (**Extended Data Fig. 3d**).

### Erythroid-specific *Got2* deletion results in anemia and elevated erythroid-restricted progenitor cells

To define the functional relevance of GOT2 within the erythroid lineage, we then crossed *Got2* floxed mice^22^ to mice expressing a tamoxifen-inducible erythroid-specific Cre recombinase, *Gata1-CreERT2*^23^ (hereafter referred to as *Got2* cKO). Following initiation of tamoxifen diet (**Extended Data Fig. 4a**), *Got2* cKO mice exhibit *Got2* deletion in *Gata1*-expressing megakaryocytic-erythrocytic progenitors (MEPs) and subsequent erythroid progenitors and precursors, as shown previously.^39^ We confirmed efficient *Got2* deletion in sorted erythroid progenitors (**Extended Data Fig. 4b**). Since the *Gata1-CreERT2* allele was previously shown to have no impact on erythropoiesis on its own^23^, *Got2^f/f^* mice were used as controls. *Got2* cKO mice exhibited significant microcytic anemia 1 month post-tamoxifen administration, with a ∼35% reduction in RBC count, as well as significant splenomegaly (**Fig. 2b, Extended Data Fig. 4c,d**). The anemia became more severe 2 months post-tamoxifen, with a ∼55% reduction in RBC count. Interestingly, the anemia became macrocytic at the later time point (**Extended Data Fig. 4e**), which could be due to increased reticulocyte count. Indeed, the reticulocyte count appeared to be increased at 2 months, but not 1 month, post-tamoxifen administration (**Extended Data Fig. 4c,e**).

*Got2* deletion from erythroid progenitors and precursors would be expected to decrease aspartate levels. Due to technical limitations of collecting sufficient progenitor and precursor cells for reliable metabolomics, we turned to the circulating RBC to understand the metabolic changes occurring with *Got2* deletion. Circulating RBCs do not have mitochondria, and thus GOT2 is absent from both control and *Got2* cKO mature RBCs. With this in mind, we observed a nonsignificant downward trend in aspartate levels in RBCs harvested from *Got2* KO mice (**Fig. 2c**) with no other notable metabolic changes identified (**Extended Data Fig. 4f-h**). This indicates that, while fewer, *Got2* cKO RBCs that are able to develop appear metabolically identical to control.

To characterize the stages of erythroid differentiation that require GOT2, we harvested and stained bone marrow cells using conventional cell surface markers^40,41^ to distinguish pre-CFU-E, colony-forming unit-erythroid (CFU-E), and terminally mature erythroid precursors (**Extended Figure Fig. 5a-c**). The frequency of late burst-forming unit-erythroid (BFU-E) and early CFU-E, collectively defined as pre-CFU-E, was significantly increased in *Got2* KO mice (**Fig. 2d**). Similarly, the frequency of CFU-E cells trended upwards in *Got2* cKO mice (**Fig. 2d**). In contrast, the absolute number of the more differentiated TER119+ erythroblasts was significantly reduced in *Got2* cKO mice (**Fig. 2d**).^42^ Specifically, the absolute numbers of polychromatic erythroblasts (PolyEbs) and orthochromatic erythroblasts (OrthoEbs) were notably reduced in *Got2* KO mice (**Extended Data Fig. 4i**). Taken together, these data suggest a block in erythroid differentiation resulting from GOT2 deficiency. This result appeared to be supported when we examined existing murine mRNA expression databases for *Got2* expression in the erythroid lineage. *Got2* is highly expressed in mouse erythroid progenitors, but its expression decreases rapidly during erythroblast maturation^17^ (**Fig. 2e**).

Curiously, the platelet count was elevated in *Got2* cKO mice (**Extended Data Fig. 4c,e**). However, the percentage of bone marrow megakaryocyte progenitors was unchanged (**Extended Data Fig. 4j**), suggesting that the increase in platelet count was not due to competition between erythroid and megakaryocyte differentiation in the setting of GOT2 deficiency in MEP cells.

### Erythroid-specific *Got1* deletion results in anemia and elevated erythroid-restricted progenitor cells

Aspartate is made by GOT2 and actively pumped out of the mitochondria by the glutamate-aspartate antiporter using energy from the mitochondrial proton gradient (**Fig. 2a**).^43^ In the cytosol, catabolism of aspartate by GOT1 is driven by substrate availability. Previous work in HSCs demonstrated that *Got2* and *Got1* play opposing roles in HSC maintenance, consistent with their functions in making and breaking aspartate, respectively.^24^ Namely, *Got2* deletion decreases aspartate and HSC regeneration, while *Got1* deletion increases aspartate and HSC regeneration. The authors showed that aspartate is required for purine and asparagine biosynthesis. This led us to hypothesize that erythroid-specific *Got1* deletion would similarly increase aspartate levels in erythroid progenitors and result in elevated blood counts reflective of increased erythropoiesis. Therefore, we generated *Gata1-CreERT2;*Got1^Δ/Δ^ (hereafter referred to as *Got1* cKO) mice, and confirmed *Got1* deletion in isolated erythroid progenitors (**Extended Data Fig. 6a,b**). We similarly showed reduced GOT1 protein levels in circulating RBCs from *Gata1-CreERT2;*Got1^Δ/Δ^ mice (**Extended Data Fig. 6c**).

In contrast with previous work in HSCs, *Got1* deletion resulted in normocytic anemia 1 month post-tamoxifen administration, with a ∼20% reduction in RBC counts, associated with significant splenomegaly (**Fig. 2f, Extended Data Fig. 6d,e**). The degree of anemia appeared unchanged at 2 months, but it became macrocytic (**Extended Data Fig. 6f**), which could be due to increased reticulocyte count.

Metabolomic analysis of circulating RBCs confirmed significantly elevated aspartate levels in *Got1* cKO mice compared to control (**Fig. 2g**), with several other metabolite changes (Extended Data Fig. 6g-i). Similar to *Got2* cKO mice, *Got1* cKO mice also exhibited an increase in pre-CFU-E cells and a decrease in terminally differentiated TER119+ erythroblasts (**Fig. 2h**). Specifically, the absolute numbers of basophilic erythroblasts (BasoEbs), PolyEbs and OrthoEbs were reduced in *Got1* KO mice (**Extended Data Fig. 6j**). The similar necessity of the GOTs in erythropoiesis seemed even more apparent when comparing the analogous murine mRNA expression of *Got1* during erythroid differentiation (**Fig. 2i**) to *Got2*. Notably, *Got1* cKO mice did not exhibit thrombocytosis (**Extended Data Fig. 6d,f**).

### Erythroid-specific double *Got1*/*Got2* deletion results in anemia and elevated erythroid-restricted progenitor cells

To assess potential compensation between GOT1 and GOT2 by reversal of their canonical reactions, we next generated *Gata1-CreERT2; Got1^Δ/Δ^Got2^Δ/Δ^* mice (hereafter referred to as *Got1/2* cDKO). Following tamoxifen administration, *Got1/2* cDKO mice exhibited similar trends in blood counts as *Got2* KO mice. One month post-tamoxifen, *Got1/2* cDKO mice demonstrated microcytic anemia, with a 29% reduction in RBC count, and significant splenomegaly (**Fig. 2J, S7A-B**). The anemia became more severe and macrocytic 2 months post-tamoxifen, with a ∼52% reduction in RBC count (**Fig. S7C**). *Got1/2* cDKO mice also exhibited an increase in pre-CFU-E, a decrease in terminally differentiated TER119+ erythroblasts (**Fig. 2L, S7G**), and increased platelet counts (**Fig. S7A, S7C**).

Metabolomic analysis showed significantly elevated aspartate levels in RBCs harvested from *Got1/2* cDKO mice compared to control RBCs (**Fig. 2K**), with other metabolite changes resembling the Got1 cKO (**Fig. S7D-F**). This is likely the consequence of cells retaining cytosolic GOT1 expression, whereas GOT2 is lost when mitochondria are removed during the final stages of blood development. While the metabolome of RBCs is different than that of erythroid progenitors and precursors, the aspartate level is increased in *Got1/2* cDKO and *Got1* cKO RBCs and decreased in *Got2* cKO. Yet, both *Got1/2* DKO and *Got2* single KO mice exhibited similar levels of anemia. These data are further suggestive that defective erythropoiesis results from a mechanism separate from regulation of aspartate levels. Collectively, these data suggest that there is not Got isoform compensatory activity in erythroid cells.

### Myeloid-specific double Got1/2 deletion does not cause changes in myeloid cell counts

To further explore whether GOT activity is uniquely important within the erythroid lineage, we used a conditional knockout approach in the myeloid lineage. We bred *Got1* and *Got2* floxed mice to those expressing the *LysM-Cre*, which is myeloid-specific^44,45^, generating *LysM-Cre;Got1^Δ/Δ^Got2^Δ/Δ^* (*LysM* cDKO) mice. Confirmation of *Got1* and *Got2* depletion efficiency was achieved through evaluating protein levels in bone marrow-derived macrophages (**Fig. S7H**). Within the blood, we found no significant changes in the number of circulating myeloid cells (**Fig. S7I**). This is in alignment with previous research showing that *Got1* is dispensable for macrophage function^46^. These studies suggest that GOT activity is dispensable for the generation of some blood lineages, such as myeloid cells, while remaining critical for the generation of others, such as erythroid cells.

### *Got2* deletion does not appear to impair erythropoiesis through NADH reductive stress

While the role of aspartate has been highlighted in regenerating HSCs, with opposite findings in *Got1* and *Got2* deleted HSCs^24^, the observation that *Got1* and *Got2* deletion in erythroid cells results in a similar anemia phenotype suggested an alternative function for GOT1 and GOT2 in erythroid cells. While *Got1* and *Got2* deficiency lead to opposite effects on aspartate levels, both are expected to disrupt the ability of the MAS to transfer reducing equivalents, leading to an elevation in the cytosolic NADH/NAD+ ratio, defined as NADH reductive stress. This subsequently leads to metabolic dysfunction, prominently including inhibition of glycolysis and oxidative phosphorylation.^22^

To test this, we focused on MDH1, another enzyme in the MAS, as it has previously been shown that MDH1-mediated MAS is important f or the activity of fetal liver HSCs. Moreover, similar to the expression patterns of *Got1* and *Got2* during blood cell differentiation (**Fig. 2e,i**), *Mdh1* is highly expressed in mouse erythroid progenitors and decreases rapidly during erythroblast maturation (**Fig. 3a**). Thus, we hypothesized that *Mdh1* deletion would phenocopy *Got* loss, similarly leading to a dysfunctional MAS and NADH reductive stress. We generated and crossed an *Mdh1* floxed allele with *Gata1-CreERT2* to create the *Mdh1* cKO model. Following the same protocol as described for the *Got* cKO models, we did not observe significant changes in blood counts or spleen size in *Mdh1* cKO mice after 1 month of tamoxifen (**Fig. 3b**). MDH1 knockout was verified by analyzing MDH1 protein levels in circulating RBCs of *Mdh1* KO mice (**Fig. 3c**). These data indicate that the MAS control of NADH reductive stress is not the mechanism behind GOT-loss driven anemia.

**Fig. 3:**
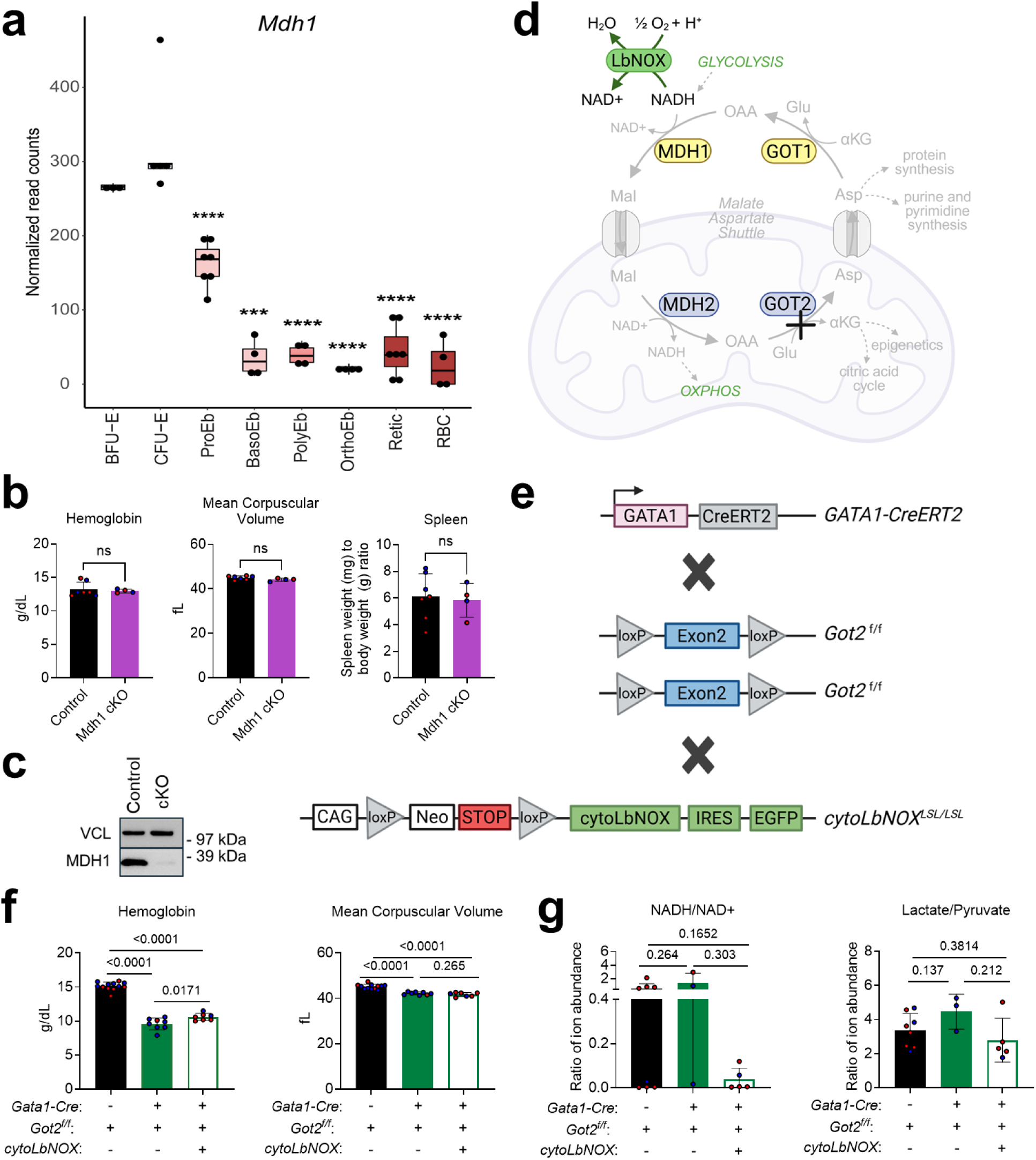
Maintenance of NADH reductive stress through the MAS is dispensable for blood cell development. a, The relative mRNA expression of *Mdh1* during mouse erythropoiesis, shown as normalized mRNA read counts by RNA-seq. **b,** Hemoglobin and mean corpuscular volume of control (n=7) or *Mdh1* cKO (n=4) mice. Control mice are *Mdh1^f/f^*or *Gata1-Cre;Mdh1^Δ/+^.* ns: nonsignificant. **c,** Western blot of MDH1 and VINCULIN loading control in circulating RBCs of control and *Gata1-Cre;Mdh1^Δ/Δ^* (cKO) mice, showing that MDH1 is depleted. **d,** Cytoplasmic *Lb*NOX (cyto*Lb*NOX) oxidizes NADH by transferring its electrons to oxygen, yielding NAD+ and water, and thereby relieving reductive stress and allowing glycolysis and OXPHOS to resume. **e,** *cytoLbNOX^LSL/LSL^* mice are characterized by a Lox-Stop-Lox (LSL) cassette upstream of c*Lb*NOX and an internal ribosomal entry site–linked enhanced GFP. To achieve erythroid-specific knock out of *Got2* and expression of cyto*Lb*NOX, we crossed the *Got2* cKO mice with the *cytoLbNOX^LSL/LSL^* mice, resulting in *Gata1-CreERT2;Got2^Δ/Δ^cLbNOX ^LSL/+^*(*Got2* cKO;c*Lb*NOX) mice and control *Got2^fl/fl^;cytoLbNOX^LSL/+^*mice. **f,** Hemoglobin and mean corpuscular volume of *Got2^f/f^*(black bar, n=12), and *Got2* cKO (green bar, n=7), *Got2 KO;*cyto*Lb*NOX (green outlined bar, n=7). **g,** Relative ratios in the indicated metabolites in the *Got2^f/f^* (black bar, n=8), *Got2* cKO (green bar, n=3), and *Got2* cKO;cyto*Lb*NOX (green outlined bar, n=5). Red and blue circles represent female and male mice, respectively. The statistical significance test for all graphs was done with an unpaired two-tailed t-test with a significance threshold level of 0.05.

To further validate this finding, we applied an orthogonal approach. The reductive stress from MAS inhibition can be relieved if the cell has access to electron acceptors. One such electron acceptor is oxygen, and *Lactobacillus brevis* NADH oxidase (*Lb*NOX) is a bacterial enzyme that transfers the reducing power in NADH onto oxygen to make NAD+ and water^47^ (**Fig. 3d**), thereby ameliorating NADH reductive stress. Previously, we demonstrated that cytoplasmic *Lb*NOX can rescue NADH reductive stress in *GOT2* KO pancreatic cancer cells.^22^ To determine if NADH reductive stress is responsible for anemia in the setting of erythroid-specific *Got2* deletion, *Got2* cKO mice were engineered to express cytoplasmic *Lb*NOX (cyto*Lb*NOX), called *Got2* cKO;cyto*Lb*NOX mice (**Fig. 3e**). Notably, cyto*Lb*NOX did not rescue the anemia in *Got2* cKO;cytoLbNOX mice (**Fig. 3f**). To assess the metabolic activity of cyto*Lb*NOX, we measured NADH/NAD+ ratios and lactate/pyruvate ratios in circulating RBCs. In both cases, there was a downward trend in these ratios when cyto*Lb*NOX was present, suggesting the rescue of NADH reductive stress (**Fig S3g**). This data further suggests that the mechanism of anemia from GOT-loss is dependent on one of their roles outside of the MAS

### *GOT1* and *GOT2* deletion decreases RBC formation and leads to epigenetic changes in histone methylation

Next, we utilized human erythroid cells to define the role of GOT1 and GOT2, given the known differences between murine and human erythropoiesis^48–50^. We first assessed whether GOT1 and GOT2 deletion would similarly impair human erythroid development utilizing primary human CD34+ hematopoietic stem and progenitor cells (HSPCs) directed to undergo erythroid differentiation *in vitro* (**Fig. 4a**). To replicate the developmental stage at which *Gata1-CreERT2* is activated in mice, we performed electroporation at day 2 of CD34+ cell differentiation (**Fig. 4a**). Deletion efficiency was confirmed by Western blot (**Extended Data Fig. 8a**), and metabolomic analysis of day 14 erythroid cells showed an increase in aspartate levels in GOT1 deleted, but not GOT2 deleted cells (**Extended Data Fig. 8b**). We also observed signs of NADH reductive stress in both GOT1 or GOT2 deleted cells (**Extended Data Fig. 8c,d**).

**Fig. 4:**
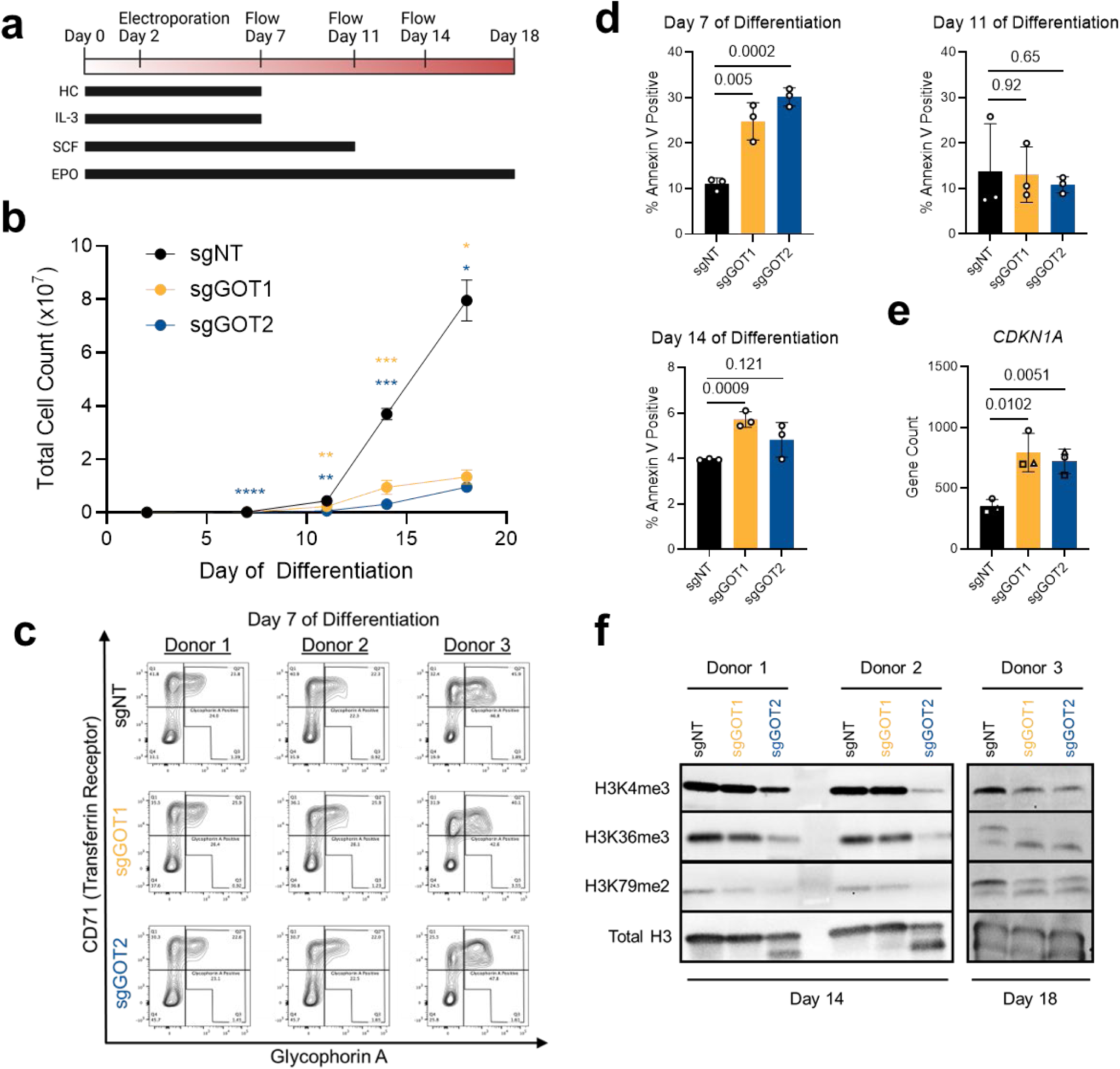
*GOT* knockout impairs human erythroid progenitor cell proliferation and survival. **a,** Schematic illustrating the 18-day differentiation strategy of human CD34+ hematopoietic stem and progenitor cells (HSPCs) into erythroid cells *in vitro*. Deletion of *GOT1* and *GOT2* was done at day 2 using Cas9/sgRNA ribonucleoprotein electroporation. HC: hydrocortisone; SCF: stem cell factor. **b,** Total counts of erythroid cells differentiated from CD34+ HSPCs following transfection with a nontargeting sgRNA (sgNT), a *GOT1*-targeting sgRNA (sgGOT1), or a *GOT2-*targteing sgRNA (sgGOT2). Similar experiments in 2 other donors exhibit the same trends. * p<0.05; ** p<0.01; *** p<0.001; **** p<0.0001. **c,** Erythroid differentiation of CD34+ HSPCs assessed by flow cytometry at day 7 of differentiation. Live, singlet cells were analyzed for Glycophorin A and CD71 expression. **d,** Proportion of singlet cells that are positive for Annexin V at the indicated timepoints. **e,** Count per million of *CDKN1A* mRNA transcripts from bulk RNA sequencing analysis of HSPC-derived erythroid cells at day 5 of differentiation. Experiment was done using 3 independent HSPC donors. Closed circle data point is donor 1, open circle is donor 2, and triangle is donor 3. **f,** Western blot of H3K4me3, H3K36me3, H3K79me2 and total H3 loading control in HSPC-derived erythroid cells at day 14 and day 18 of differentiation. The statistical significance test for all graphs was done with an unpaired two-tailed t-test with a significance threshold level of 0.05.

*GOT1* or *GOT2* knockout resulted in reduced cell counts in culture (**Fig. 4B**), without impairment in erythroid differentiation at day 7, as demonstrated by glycophorin A and CD71 (transferrin receptor) staining (**Fig. 4c**). These results reflect those from our mouse studies, with the most significant impact on ultimate RBC number, while the cells that manage to transit beyond the pre-CFU-E differentiation block look otherwise normal. Further, the lack of differentiation defect in this human system has been seen in previous work, where downregulation of GOT1 in CD34+ HSPCs with shRNA yielded no differentiation defect *in vitro*.^51^ To understand the reduction in cell numbers, we looked at the apoptotic marker Annexin V and found it to be increased at day 7 of differentiation following *GOT1* and *GOT2* deletion (**Fig. 4d**). Additionally, using bulk RNA sequencing, we surveyed gene expression changes in erythroid cells derived from 3 independent HSPC donors at day 5 of differentiation. The top upregulated gene was *CDKN1A*, which plays a role in arresting the cell cycle (**Fig. e**). Gene set enrichment analysis further confirmed an upregulation in genes associated with apoptosis in *GOT1* or *GOT2* depleted erythroid cells at day 5 (**Extended Data Fig. 9a**). These data support a critical role for GOT1 and GOT2 in early erythropoiesis, both in mice and in humans.

We previously concluded from our murine models that the maintenance of aspartate levels and the MAS are not the key functions of the GOT enzymes. We next hypothesized that disruption of αKG metabolism might dysregulate key epigenetic programs required for erythropoiesis. Strikingly, we observed decreased methylation of several activating histone modifications, H3K4me3, H3K36me3, and H3K79me2, in *GOT2* deleted cells at day 14 and in both *GOT1* and in *GOT2* deleted cells at day 18 (**Fig. 4f, Extended Data Fig. 9b**). Di-and trimethylation of H3K4, H3K36, and H3K79 are associated with transcriptional activation, so their reduction suggests enhanced gene repression. In culture, we previously demonstrated that GOT knockout in pancreatic cancer cells slows their proliferation.^22,52^ However, the reduction in methylation marks was not observed in the pancreatic cancer cells following GOT loss (**Extended Data Fig. 9c**), suggesting that these may be erythroid-specific epigenetic changes.

## Discussion

In this report, we describe increased aspartate levels in multiple, independent models of erythropoiesis. This observation led us to speculate that aspartate availability may be necessary for efficient erythropoiesis.

However, manipulating aspartate levels in either direction through *Got-*deletion resulted in similar anemic outcomes. This intriguing observation, which is in marked contrast to what is seen in HSCs^24^, led us to explore the mechanism by which the GOTs are required for efficient erythropoiesis.

Using mouse models with temporal *Got* deletion, we discovered that the GOTs are crucial in early erythroid progenitors, consistent with their gene expression patterns. Moreover, we show that the severe anemic phenotype resulting from erythroid *Got2* KO leads to microcytic anemia. The GOTs are PLP-dependent enzymes, and PLP is the active form of vitamin B_6_. It is well-established that vitamin B_6_ deficiency leads to microcytic anemia.^53–56^ Thus far, it is assumed that this is because the rate-limiting enzyme of heme synthesis, δ-aminolevulinic acid synthase (ALAS), is a PLP-dependent enzyme and vitamin B_6_ deficiency decreases ALAS activity.^53–56^ However, ALAS is not expressed until terminal erythroid differentiation, when erythroblasts are ramping up hemoglobin synthesis.^54^ Thus, we propose that vitamin B_6_ deficiency also causes microcytic anemia due to inefficient early erythropoiesis; specifically, due to the inability of the PLP-dependent GOTs to function at this stage of blood development. Vitamin B_6_ deficiency has already been shown to dampen GOT2 activity in the setting of acute myeloid leukemia.^57^

Our discovery that the GOTs are crucial during early erythropoiesis aligns with other related studies in the field. For example, a CRISPR screen aimed at defining the genes required for terminal erythroid differentiation did not identify the GOTs as crucial genes in terminal erythroid maturation.^58^ Additionally, previous work has shown that aminooxy acetic acid (AOA), an inhibitor of PLP-dependent aminotransferases including GOT1 and GOT2, inhibits early erythroid differentiation of primary human HSPCs.^15,18^

The GOTs have canonical roles within the MAS, and thus a plausible explanation for their necessity is that deletion of either *Got* would lead to disruption of the MAS and, consequently, NADH reductive stress. This mechanism seemed even more likely as all components of the MAS showed a similar gene expression pattern, being highly expressed in mouse erythroid progenitors and decreasing during erythroblast maturation during terminal differentiation.^17^ To test this hypothesis, we deleted another enzyme within the MAS, MDH1. Similar to *Got*-deletion, *Mdh1*-deletion disrupts the MAS and leads to NADH reductive stress. In contrast to our hypothesis, *Mdh1*-deletion did not cause anemia. In further validation, alleviation of NADH reductive stress from *Got2*-deletion through ectopic expression of cytoplasmic *Lb*NOX also did not rescue the anemic phenotype.

The GOT enzymes are reversible, being driven by substrate availability. Double Got knockout addressed the possibility of compensatory activity of these enzymes; however, this does not explain why the single knockouts have the same phenotype, with Got1 resulting in a more modest impact on anemia. We propose that since GOT enzymes catalyze substrate-driven reactions, aspartate accumulation resulting from Got1 deletion may mimic reduced GOT2 activity through product-mediated feedback inhibition. Increased aspartate levels following Got1 loss could impair the ability of GOT2 to generate both aspartate and αKG. This would explain both the less severe Got1 phenotype, and why the double knockout phenocopies Got2 knockout.

Using multiple, orthogonal mouse and human models, we demonstrate that GOT regulation of NAD(H) and aspartate synthesis are not the molecular mediators of defective blood cell production. Instead, our data analyzing histone marks in human GOT deleted cells implicates αKG production. This is consistent with prior studies showing that AOA-mediated inhibition of early erythroid differentiation in primary human HSPCs could not be rescued by nucleosides, which depend on aspartate availability, but could be rescued by cell-permeable dimethyl αKG^15,18^. αKG is a co-factor for histone and DNA demethylases, and its availability directly impacts the epigenetic code and thus gene expression. During terminal erythropoiesis, heme biosynthesis imposes a substantial αKG demand.^15^ We propose that in early erythroid progenitors, this metabolic consumption of αKG is sensed by chromatin, establishing a metabolic–epigenetic checkpoint that ensures differentiation proceeds only when αKG availability can support both heme production and the associated epigenetic remodeling.

Notably, elevated αKG has been reported to reduce specific histone methylation marks via increased demethylase activity, but it can also decrease certain methyl marks through compensatory αKG-independent demethylase mechanisms. This speculation notwithstanding, it is possible that αKG may be playing multiple, non-mutually exclusive roles, including regulating gene expression and fueling the citric acid cycle. Future studies will be needed to further elucidate the mechanisms linking GOT activity, αKG metabolism, and epigenetic regulation during erythropoiesis.

### Limitations

This work was technically limited by the ability to collect sufficient progenitor and precursor cells for reliable metabolomics measurements, which led us to utilize the circulating RBC for preliminary explorations. While sufficiently abundant, the RBC state is a functionally separate cell state that has no organelles (including mitochondria and thus GOT2) and a uniquely different cellular metabolism. As such, we were only able to draw corollary conclusions about the metabolic state. Future studies assessing metabolism in this highly limiting population will be valuable to ascertain more proximal metabolic changes. This work was also limited by differences in biology between mice and human erythropoiesis, though we attempted to address this shortcoming by using the CD34+ primary human cell differentiation model to assess translatability. Further to this point, the variability within human donors led to challenges in identifying universally significant gene expression changes across all donors.

## Methods

### Animals and Procedures

Mouse experiments were conducted under the guidelines of the Office of Laboratory Animal Welfare and approved by the Institutional Animal Care and Use Committees at the University of Michigan under protocol PRO00010606 and PRO00011805. All mice were kept in ventilated racks in a pathogen-free animal facility with a 12 h light/12 h dark cycle, 30–70% humidity, and 68–74°F temperatures. All mice were maintained in the animal facility in the Rogel Cancer Center at the University of Michigan and were overseen by the Unit for Laboratory Animal Medicine (ULAM). Male and female mice were used for each experiment.

#### Phenylhydrazaline Treatment

Mice were treated for two consecutive days with phenylhydrazine (Phz, Sigma-Aldrich, St. Louis, MO) at 60 mg/kg of body weight via intraperitoneal injection. Blood was collected three days after the second injection.

#### Humanized sickle cell disease (SCD) Mouse Model

Mice were utilized that had their endogenous α-and β-globin genes replaced by fragments of the human α-and sickle β^S^-globin genes^59^. Heterozygote mating of these mice produces homozygous SCD mice, which were then bred and maintained as described previously^59^. HPLC hemoglobin electrophoresis was performed on all mice to confirm sickle cell genotype.

#### Bone Marrow Transplantation (BMT)

For BMT, mice received 12 Gy of total body irradiation, which was achieved by administering one dose of 6 Gy, a 4 hour recovery, and then administering a second dose of 6 Gy. Bone marrow transplant was performed the day after irradiation and engraftment of the donor marrow was near 100%. All mice were sacrificed 3 months following BMT.

#### Peripheral Blood Analysis

For complete blood counts (CBCs), whole blood was collected into K3 EDTA anticoagulant tubes (Microvette 20.1288.100) and CBCs were run on a Heska Element HT5 (Heska Corporation, Loveland, CO) automated veterinary hematology analyzer. Assays were performed within the ULAM In Vivo Animal Core pathology laboratory at the University of Michigan. Quality control was performed daily using manufacturer-provided reagents, and the laboratory is a participant in an external independent quarterly quality assurance program (Veterinary Laboratory Association Quality Assurance Program).

### Mouse strains

*Ubiquitin-CreERT2;Got2^Δ/Δ^.* Transgenic mice with a tamoxifen inducible Cre-ERT2 expressed from the human ubiquitin C (UBC) promoter were used.^38^ UBC-Cre-ERT2 mice were bred with the *Got2* floxed allele^22^, generating the UBC-Cre-ERT2;*Got2^Δ/Δ^* mouse model. The primers used for genotyping the tails of 3-week old mice are listed in Supplementary Table 2. Eight to twelve week old mice were administered 5 mg tamoxifen by oral gavage daily for five days. Animals were then fed 400 mg tamoxifen citrate per kg chow (Envigo TD.130860) for 3 weeks to achieve whole-body Cre recombination to delete *Got2*.

*LysM-Cre;Got1^Δ/Δ^Got2^Δ/Δ^*. LysM-Cre mice^45^ (Jackson Lab strain #004781) on a pure C57BL/6J background were crossed with *Got1^fl/fl^* mice^60^ and *Got2^fl/fl^* mice^22^ that were also on a pure C57BL/6J background. *LysM-Cre;Got^Δ/Δ^* (either male or female) mice were crossed with *Got^fl/fl^* mice to generate an F1 generation of all *Got^fl/fl^* pups with or without LysM-Cre expression. The primers used for genotyping the tails of 3-week old mice are listed in Supplementary Table 2.

*Gata1-CreERT2;Got1^Δ/Δ^ and Gata1-CreERT2;Got2^Δ/Δ^*. Transgenic mice with a tamoxifen-inducible erythroid lineage-specific Cre (*Gata1-CreERT2*)^23^ were crossed with *Got1* floxed^60^ or *Got2* floxed lines^22^ to create

*Gata1-CreERT2;Got1^Δ/Δ^* and *Gata1-CreERT2;Got2^Δ/Δ^*mice. Since the presence of the *Gata1-CreERT2* allele does not result in an erythroid phenotype on its own^23^, *Got1^fl/fl^*and *Got2^fl/lf^* mice were used as controls. The primers used for genotyping the tails of 3-week old mice are listed in Supplementary Table 2. Gata1-CreERT2 activity was induced in five to ten-week-old mice following administration of 400 mg tamoxifen citrate per kg chow (Envigo TD.130860) for 1-2 months.

*Gata1-CreERT2;Got2^Δ/Δ^cytoLbNOX^LSL/+^*. *cytoLbNOX^LSL/LSL^* mice contain a Lox-Stop-Lox (LSL) cassette upstream of a cytosolic *Lb*NOX (cyto*Lb*NOX^LSL/LSL^) gene and an internal ribosomal entry site–linked enhanced GFP^61^. These mice were crossed to either *Got2^fl/fl^* mice or *Gata1-CreERT2;Got2^Δ/Δ^* mice to obtain *Got2^fl/fl^cytoLbNOX^LSL/+^*or *Gata1-CreERT2;Got2^Δ/Δ^cytoLbNOX ^LSL/+^* mice. The primers used for genotyping the tails of 3-week old mice are listed in Supplementary Table 2. Two separate genotyping reactions were done: the first to test for the presence of the LSL allele and the second to check for the presence of the allele that has the stop cassette removed (cyto*Lb*NOX expressed). Gata1-CreERT2 activity was induced in five to ten-week-old mice following administration of 400 mg tamoxifen citrate per kg chow (Envigo TD.130860) for 1-2 months.

Gat*a1-CreERT2; Mdh1^Δ/Δ^.* Mice containing loxP sites flanking exon 2 of the Mdh1 gene were generated by Ozgene. These mice were crossed to the Gata1-Cre model^23^ to create *Gata1-CreERT2;Mdh1^Δ/Δ^* mice. Since the presence of the *Gata1-CreERT2* allele does not result in an erythroid phenotype on its own^23^, *Mdh1^fl/fl^*mice were used as a control. The primers used for genotyping the tails of 3-week old mice are listed in Supplementary Table 2. Gata1-CreERT2 activity was induced in five to ten-week-old mice following administration of 400 mg tamoxifen citrate per kg chow (Envigo TD.130860) for 1-2 months.

### Cell culture

#### MEL and K562 Differentiation

MEL cells were cultured in DMEM (Gibco 11965092) supplemented with 10% FBS, 1% GlutaMAX (Gibco 35050061), 1% non-essential amino acids (Gibco 11140050), and 1% penicillin-streptomycin (Gibco 15140122). K562 cells were cultured in RPMI-1640 (Gibco 11875093) supplemented with 10% FBS, 1% GlutaMAX and 1% penicillin-streptomycin. For differentiation, cells were plated at 25,000 cells/mL and treated the day after with 2% DMSO (Sigma D2650) for MEL cells or 1.5mM sodium butyrate (Sigma B5887) for K562 cells. After 48 hours, cells were diluted with the same volume of media they were originally plated in. This additional media was supplemented with 2% DMSO or 1.5mM sodium butyrate. Cells were collected 48 hours after this second treatment. Cells were maintained at 37°C in 5% CO_2_.

#### CD34+ Differentiation Assays

Human peripheral blood-derived CD34^+^ HSPCs were purchased from the Fred Hutchinson Cancer Research Center and were cultured and differentiated into mature erythroid cells using a 3-phase culture system reported previously.^62^ In brief, cells were cultured in an erythroid differentiation medium (EDM), which consisted of RPMI 1640 (Thermo Fisher 11875093) supplemented with 5% virus-inactivated human plasma AB (Rhode Island Blood Center X00004), 1% glutamine (Thermo Fisher/Gibco 35050-061), 1% Pen-Strep (Thermo Fisher/Gibco 15140-122), 330mg/L human holo-transferrin (Gemini Bio 800-131P), 10mg/L recombinant human insulin (Sigma 91077C), 2 U/L heparin (Sigma H3149), and 3U/L erythropoietin (Amgen).

The expansion procedure comprised 3 phases. In the first phase (day 0 to day 7), 10^4^/mL CD34+ cells were cultured in EDM described above supplemented with 10^-6^ M hydrocortisone (StemCell Tech 74142),100 ng/mL SCF (R&D 255-SC-050) and 5 ng/mL IL-3 (R&D 203-IL). In the second phase (day 7 to day 11), cells were resuspended at 10^5^/mL in EDM supplemented with 100 ng/mL SCF. In the third phase (day 11 to day 18), cells were resuspended in EDM supplemented with an additional 660mg/L holo-transferrin.

Cells were maintained at a concentration of 10^4^-10^6^ cells/mL from days 1 to 7, 10^5^-10^6^ cells/mL from days 7 to 11, and 2.5×10^5^-10^6^ cells/mL between days 11 to 18. All cultures were maintained at 37°C in 5% CO2 and 21% O_2_.

#### Generation of CRISPR/Cas9 knockout CD34+ cells

Human CD34+ HSPCs were electroporated on day 2 of differentiation using a protocol reported previously.^63^ In brief, cells were first washed two to three times in PBS to remove residual RNAses present in the culture media. Meanwhile, a master mix of Cas9/sgRNA ribonucleoprotein was prepared by combining 0.5μL of 100μM sgRNA in ultrapure water with 1.2μL of 10μg/μL Alt-R S.p. HiFi Cas9 Nuclease V3 (IDT, 1081061). This was gently swirled with a pipette tip and the mix was incubated at room temperature for 10-30 minutes. 1×10^5^ to 2×10^5^ cells were resuspended in 16.4 μL P3 Primary Cell Nucleofector Solution (Lonza, V4XP-3032), 3.6 μL Supplement 1 (Lonza, V4XP-3032) and 1uL of 100uM IDT electroporation enhancer reagent (IDT #1075916). The 21μL cell suspension was then combined with the complexed Cas9 ribonucleoprotein master mix, gently mixed four times, and the 22.7μL was transferred to an electroporation cuvette. Cells were electroporated using the DZ-100 program in a 4D-Nucleofector X Unit (20 μL cuvettes). Immediately after electroporation, 80 μL of prewarmed media was added to the electroporation cuvette and cells were gently mixed before plating at a density of 100,000 cells/mL in prewarmed media. sgNT (Non-targeting): CGCUUCCGCGGCCCGUUCAA; sgGOT1: GAUAGGCUGAGUCAAAGAAG; sgGOT2: GUGCGCCACUUCAUCGAACA

#### Non-Nuclear Western Blotting

Suspension cells were collected and spun down at 350g at 4°C for 5 minutes. The resulting supernatant was aspirated off, PBS was added to wash the cell pellet, the pellet was spun again at 350g at 4°C for 5 minutes, and the PBS was aspirated off. The washed cell pellet was resuspended in ∼300-500mL of a lysis buffer containing RIPA (Sigma Aldrich), phosphatase inhibitor (Sigma Aldrich), and proteinase inhibitor (Sigma Aldrich). This was left on ice for 10 minutes, the pellet vortexed, and then spun for 10 minutes at 4°C at maximum centrifugal speed to pellet the cellular debris. The resulting supernatant containing the extracted proteins was moved to a new tube.

Protein concentrations were determined using a colorimetric bicinchoninic acid assay (Thermo Fisher), according to manufacturer’s instructions. Following quantification, the necessary volume of lysate containing 10-20 µg of protein was added to a mixture of sample buffer (Invitrogen) and reducing agent (Invitrogen) and incubated at 85°C for 10 minutes. Next, the lysate was separated on a 4-12% Bis-Tris gradient gel (Invitrogen) along with a protein ladder (Invitrogen) at 100 V until the dye reached the bottom of the gel (about 120 minutes). Then, the protein was transferred to a methanol-activated PVDF membrane (Millipore) at 20 V for 1 hour. After that, the membrane was blocked in 5% blocking reagent (Biorad) dissolved in TBS-T (TBS buffer containing 0.1% Tween-20) on a plate rocked for 1 hour. The membrane was then incubated overnight at 4°C rocking in the indicated primary antibody diluted in 5% BSA in TBS-T. The next day, the primary antibody was removed, and the membrane was washed 3 times in TBS-T rocking for 5 minutes. Then, the membrane was incubated for 1 hour rocking at room temperature in the appropriate secondary antibody diluted in TBS-T. Finally, the membrane was washed and incubated in Clarity ECL reagent (Biorad) according to manufacturer’s instructions before imaging on a Biorad Chemidoc. The following antibodies (Supplemental Table 3) were used for this study: Vinculin (Cell Signaling 13901, 1:10,000 dilution), GOT2 (Sigma, #HPA018139, 1:1000 dilution), GOT1 (Abcam, ab171939, 1:1000 dilution), and MDH1 (Abcam, ab180152, 1:6000 dilution).

#### Nuclear Western Blotting

The cells were collected as described in non-nuclear Western blotting. The cell pellet was resuspended in 1ml of hypotonic lysis buffer (containing 10 mM Tris HCl pH 8.0, 1 mM KCl, 1.5 mM MgCl2, protease inhibitor (100X stock, Sigma Aldrich, P8340) and 1X phosphatase inhibitor (100X stock, Sigma Aldrich, #P5726)) and incubated for 30 minutes on a rotator at 4°C. The pellet was then collected by centrifugation at 10,000g, 4°C for 10 minutes. It was subsequently resuspended in 400 μl of 0.4 N H_2_SO_4_ and left to incubate overnight on a rotator at 4°C. After centrifugation at 16,000g, the resulting supernatant was transferred to a new tube, and 132 μl of trichloroacetic acid was gradually added. The mixture was then incubated on ice for 30 minutes. The histone pellet was obtained by centrifugation at 16,000g, 4°C for 10 minutes and washed with cold acetone. The suspension was centrifuged at 16,000g, 4°C for 5 minutes and the acetone was removed. The histone pellet was further washed with acetone and subsequently allowed to air dry with the caps open at room temperature for 20 minutes to eliminate any remaining acetone. Finally, the dried histone pellet was resuspended in an appropriate volume of ddH2O, supplemented with protease inhibitors (Sigma), and kept on ice.

Protein concentrations were determined using a colorimetric bicinchoninic acid assay (PierceTM BCA Protein Assay #23227), according to manufacturer’s instructions. Following quantification, the necessary volume of lysate containing 0.5 µg of protein was added to a mixture of loading dye (Thermo Fisher) and reducing agent (Thermo Fisher) and incubated at 85°C for 10 minutes. Next, the lysate was separated on a 4-15% Mini-Protean TGX precast gels (Biorad #3450027) along with a protein ladder (Biorad) at 100 V until the dye reached the bottom of the gel (about 90 minutes). Then, the protein was transferred to a methanol-activated PVDF membrane (Biorad #10026933) at 12 V for 30 minutes using a Bio-Rad Trans-Blot Turbo transfer system (Biorad #1704150). After that, the membrane was blocked in 5% bovine serum albumin (Biorad) dissolved in TBS-T (TBS buffer containing 0.1% Tween-20) on a plate rocked for 1 hour. The membrane was then incubated overnight at 4°C rocking in the indicated primary antibody diluted in 5% BSA in TBS-T. The next day, the primary antibody was removed, and the membrane was washed 3 times in TBS-T rocking for 5 minutes.

Then, the membrane was incubated for 1 hour rocking at room temperature in the appropriate secondary antibody diluted in TBS-T. Finally, the membrane was washed and incubated in Pierce ECL Western Blotting Substrate (Thermo Fisher #32106) according to manufacturer’s instructions before imaging on a Thermo Fisher iBright Imaging System. The following antibodies (Supplemental Table 3) were used for this study: Total H3 (Cell Signaling Technology #3638, 1:5,000), H3K4me3 (Cell Signaling Technology #9751, 1:1,000 dilution), H3K36me3 (Activ Motif, #61021, 1:1,000 dilution), and H3K79me2 (Cell Signaling Technology #5427, 1:1,000 dilution).

#### RNA isolation and reverse transcriptase with qPCR

RNA samples were isolated using the RNEasy Micro Kit (QIAGEN, 74004) for day 5 HSPC-derived erythroid cells or RNEasy Plus Mini Kit (QIAGEN, 74134) for all other cells according to the manufacturer’s instructions. RNA purity was assessed using a NanoDrop One (ThermoFisher Scientific, ND-ONE-W). Thereafter, 1 μg of the RNA samples were reverse transcribed to cDNA using the iScript cDNA Synthesis Kit (Bio-Rad, 1708890) according to the accompanying instructions. qPCR was performed on the QuantStudio 3 Real-Time PCR System (ThermoFisher Scientific, A28131) using Power SYBR Green PCR Master Mix (ThermoFisher Scientific, 4367659). The reactions were run at 17 µl total volume consisting of 8.5 µl SYBR, 6.5 µl nuclease free water, 1 µl of cDNA, 1 µl of 10 µM forward primer, and 1 µl 10 µM reverse primer. qPCR primer sequences are listed in Supplementary Table 1. *ACTB* was used as a housekeeping gene.

### RNA-Sequencing Analysis

*<u>Bulk RNA-seq</u>.* RNA samples were submitted to Azenta Life Sciences and 150 cycle, paired-end sequencing was performed on the Illumina HiSeq platform. The raw fastq files were checked for quality using FastQC. Adapter sequences were removed, and the reads were quality trimmed using Trimmomatic^64^ with the following parameters: HEADCROP:10 ILLUMINACLIP:TruSeq3-PE.fa:2:30:10 LEADING:3 TRAILING:3

SLIDINGWINDOW:5:20 MINLEN:115. Trimmed reads were aligned to the human GRCh38.p14 reference genome from Gencode using STAR software^65^ and features were annotated and counted using RSEM software^66^ against the basic annotation GTF file from Gencode. Differential expression analysis was performed using the DESeq2 software^67^ package in R. Volcano Plots were generated using ggplot software in R. Gene Set Enrichment Analysis (GSEA) was performed using the GSEA tool^68,69^ and the Hallmark dataset from the Molecular Signatures Database (mSigDB).

*<u>scRNA-seq reanalysis</u>.* Read counts files for the GSE226078 dataset^70^ were extracted from the NCBI GEO database. The read count files were subset for erythroid lineage cells: *BFU-E*, *CFU-E*, *Pro-E*, *BasoE*, *PolyE*, *OrthoE*, *Reticulocytes*, and *RBCs*. The subsets were mined for the expression of *Got1*, *Got*2, *Mdh1, Mdh2, Slc25a11, Slc25a12,* and *Slc25a13* using custom R code. Expression plots were created using ggplot software in R.

### Flow cytometry

*<u>Murine cells.</u>* Bone marrow cells were obtained by flushing femurs and tibias with a 26G needle in phosphate-buffered saline (PBS) supplemented with 2% FBS, and cell suspensions were filtered through a 70µm cell strainer cap on a flow tube. Spleens were mechanically dissociated by crushing through a 70µm filter into a 50mL tube. Cell numbers were manually determined with a hemacytometer.

For flow cytometric analysis of **terminally differentiated erythroblasts**^4^, 3×10^6^ bone marrow cells were resuspended in 100μL PBS with 2% FBS and then stained with fluorophore-conjugated antibodies against CD11b, Gr1, and B220 along with TER119, CD71 and CD44 (refer to Supplementary Table 3) on ice for 15-30 minutes. 0.45μg of CD44 was used and 0.2ug for all remaining antibodies. 2×10^6^ cells were used for each single-color compensation and an unstained tube. Analysis was performed as previously described.^71^

For **B-lymphocytes, T-lymphocytes, and myeloid cell** analysis, 2×10^6^ bone marrow cells were resuspended in 100μL PBS with 2% FBS and then stained with fluorophore-conjugated antibodies against CD11b, Gr1, B220, CD19, and CD3e (refer to Supplementary Table 3). 0.1μg of CD3e was used and 0.05μg for all remaining antibodies. 2×10^6^ cells were used for each single-color compensation and an unstained tube.

For flow cytometric analysis of **restricted hematopoietic progenitors**, the remaining number of bone marrow cells were lineage depleted using magnetic separation (Miltenyi Biotec 130-110-470). Lineage depleted cells were then resuspended in 100μL PBS with 2% FBS and stained with 0.2μg of antibodies (excluding CD34, for which 0.8ug was used). The HSPC populations in mouse bone marrow were analyzed as previously described^40^ using the antibodies in Supplementary Table 3, including MkP cells (Kit^+^CD41^+^CD150^+^), pre CFU-E cells (Kit^+^CD41^-^CD16/32^-^CD150^-^CD105^+^) and CFU-E cells (Kit^+^CD41^-^CD16/32^-^CD150^+^CD105^+^).

*<u>Human CD34+ Cells.</u>* 1×10^5^ cells were suspended in 100 μL phosphate buffered saline (PBS) supplemented with 0.5% bovine serum albumin (BSA) and stained with 0.5µl CD71 (transferrin receptor), 0.075μL CD235a (Glycophorin A), 2µl CD233 (Band 3), and 1.5 µl CD49d (α_4_-integrin). Cells were washed once with PBS 0.5% BSA before analysis. The samples were analyzed within 1 hour after staining with a BD LSR Fortessa and FlowJo software (BD Biosciences).

### PCR genotyping of flow sorted mouse cells

Cells were sorted into PBS supplemented with 4% FBS. Tubes were spun down at 350g for 5 minutes, and the supernatant was carefully removed. 30μL of Quick Extract (Fisher NC9904870) was then added to cells and incubated in a thermocycler following manufacturer’s protocol. A 10μL PCR reaction was then set up using 7.5μL GoTaq (Promega M712), 1.5μL each of the three 10μM genotyping primers (Supplementary Table 2), and 1.5μL of the DNA solution from the thermocycler. After PCR reaction was complete, a 2% agarose gel was run at 80V for 1 hour.

### Metabolomics

*<u>Isolating Polar Metabolites from cell lines or RBCs.</u>* Metabolites were extracted from cell lines and murine RBCs with adaptations to previously reported methods.^22,26^ For cell lines, four replicates of suspension cells for each condition were collected, centrifuged at 0.2rcf for 5 minutes to pellet the cells and the media was removed. 1mL of dry-ice cold 80% was added to the cell pellet of three replicates and immediately placed on dry ice for 10 minutes. The fourth pellet was used for BCA quantification of protein, and aliquoted volumes of each of the three replicates was normalized to the protein concentrations. The normalized metabolite volumes were dried in a speedvac.

For RBCs, murine blood was collected with cardiac puncture in an Eppendorf, allowed to clot at room temperature for ∼30 minutes, and separated into serum by centrifugation for 10 minutes at 1,000xg. Serum was removed and the RBC pellet was used for metabolite extraction. 1mL of dry ice-cold 80% methanol was added to the RBC pellet and immediately placed on dry ice for 10 minutes. The mixture was then vortexed to homogenize the pellet into the 80% methanol and then centrifuged at max speed for 10 minutes at 4°C to pellet cell debris. The supernatant was collected in a new tube and any remaining 80% methanol was dried off the cell pellet using a nitrogen blower. The remaining dried pellet was then used for BCA quantification of protein, and aliquoted volumes of metabolite extractions were normalized to the protein concentrations in the cell pellet. The normalized metabolite volumes were dried in a speedvac.

*<u>LC-MS/MS Targeted Metabolomics.</u>* Samples were run on an Agilent 1290 Infinity II LC –6470 Triple Quadrupole (QqQ) tandem mass spectrometer (MS/MS) system with the following parameters^22,26^: Agilent Technologies Triple Quad 6470 LC-MS/MS system consists of the 1290 Infinity II LC Flexible Pump (Quaternary Pump), the 1290 Infinity II Multisampler, the 1290 Infinity II Multicolumn Thermostat with 6 port valves, and the 6470 triple quad mass spectrometer. Agilent Masshunter Workstation Software LC/MS Data Acquisition for 6400 Series Triple Quadrupole MS with Version B.08.02 is used for compound optimization, calibration, and data acquisition.

Solvent A is 97% water and 3% methanol 15 mM acetic acid and 10 mM tributylamine at pH of 5. Solvent C is 15 mM acetic acid and 10 mM tributylamine in methanol. Washing Solvent D is acetonitrile. LC system seal washing solvent 90% water and 10% isopropanol, needle wash solvent 75% methanol, 25% water. GC-grade Tributylamine 99% (Acris Organics), LC/MS grade acetic acid Optima (Fisher Chemical), InfinityLab Deactivator additive, ESI–L Low concentration Tuning mix (Agilent Technologies), LC-MS grade solvents of water, and acetonitrile, methanol (Millipore), isopropanol (Fisher Chemical).

An Agilent ZORBAX RRHD Extend-C18, 2.1 × 150 mm and a 1.8 μm and ZORBAX Extend Fast Guards for ultra high performance liquid chromatography (UHPLC) are used in the separation. LC gradient profile is: at 0.25 mL/min, 0–2.5 min, 100% A; 7.5 min, 80% A and 20% C; 13 min 55% A and 45% C; 20 min, 1% A and 99% C; 24 min, 1% A and 99% C; 24.05 min, 1% A and 99% D; 27 min, 1% A and 99% D; at 0.8 mL/min, 27.5–31.35 min, 1% A and 99% D; at 0.6 mL/min, 31.50 min, 1% A and 99% D; at 0.4 mL/min, 32.25–39.9 min, 100% A; and at 0.25 mL/min, 40 min, 100% A. Column temperature is kept at 35°C, samples are at 4°C, injection volume is 2 μL.

The 6470 Triple Quad MS is calibrated with the Agilent ESI-L Low concentration Tuning mix. Source parameters: gas temperature 150°C, gas flow 10 L/min, nebulizer 45 psi, sheath gas temperature 325°C, sheath gas flow 12 L/min, capillary –2000 V, and delta EMV –200 V. Dynamic multiple reaction monitoring (dMRM) scan type is used with 0.07 min peak width, acquisition time is 24 min. dMRM transitions and other parameters for each compounds are list in a separate sheets. Delta retention time of plus and minus 1 min, fragmentor of 40 eV, and cell accelerator of 5 eV are incorporated in the method.

The MassHunter Metabolomics Dynamic MRM Database and method was used for target identification. The QqQ data were pre-processed with Agilent MassHunter Workstation QqQ Quantitative Analysis Software (B0700). The metabolite abundance level in each sample was divided by the median of the control samples for proper comparisons, statistical analyses, and visualizations among metabolites. Metabolites with values >1 are higher in the experimental conditions, and metabolites with values <1 are lower in the experimental condition. The statistical significance test was done by a two-tailed t-test with a significance threshold level of 0.05. * p<0.05; ** p<0.01; *** p<0.001; **** p<0.0001. Heatmaps were generated and data clustered using Morpheus Matrix Visualization and analysis tool (https://software.broadinstitute.org/morpheus).

### Statistics

Statistics were performed using Graph Pad Prism 9 and 10. Groups of 2 were analyzed with an unpaired two-tailed student t test. All error bars represent mean with SD, all group numbers and explanation of significant values are presented within the figure legends. Experiments were repeated at least twice to verify results.

## Acknowledgements

We are deeply thankful to Dr. Amy Myers for management of experimental reagents, Ann Friedman for experimental guidance, Dr. Sean Morrison for project discussions, and Colleen Reczek for guidance with the cyto*Lb*Nox*^LSL/+^* mice. We also acknowledge the valuable services of the University of Michigan Transgenic Animal Model Core Facility.

## Author contributions

N.P., C.A.L., Y.S, and R.K. conceived the study and designed the experiments. N.P. performed most experiments, with support from other co-authors. N.P. analyzed most of the experimental data, with support from G.M. with flow cytometry data and P.S with metabolomics data. A.D.D. analyzed the RNA sequencing data. N.P., C.L., Y.S. and R.K. wrote the manuscript with assistance from all authors. All authors contributed to the evaluation, integration and discussion of the results.

## Data and materials availability

Data needed to evaluate the conclusions of the paper are present in the paper and/or the supplementary materials.

---

**Extended Data Fig 1:**
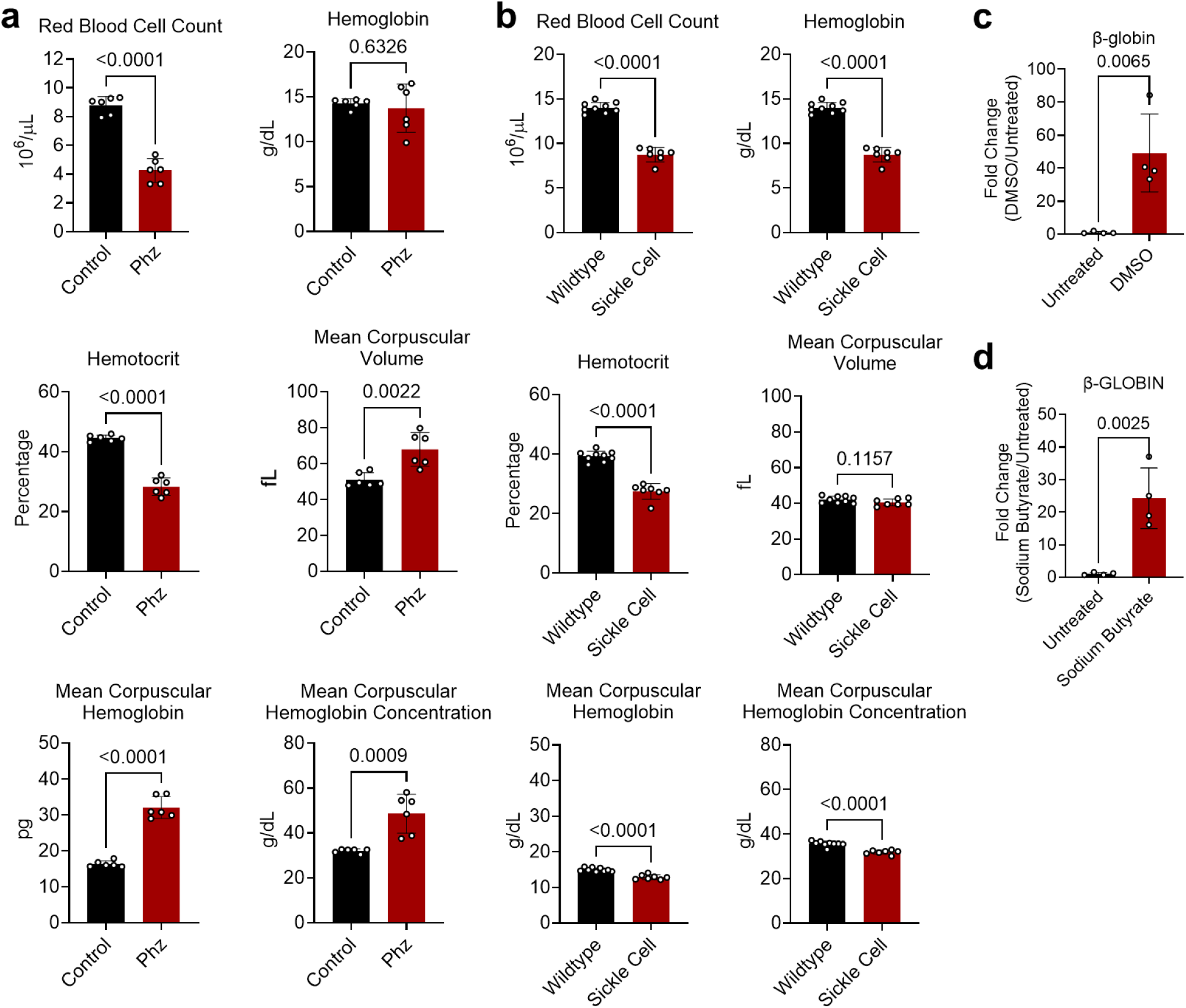
Characterization of erythropoiesis models shows decreased RBC counts in the two *in vivo* models and increased β-globin gene expression in the two *in vitro* models. a, Complete blood count analysis from control mice treated with saline (n=6) and mice treated with phenylhydrazine (Phz, n=6). b, Complete blood count analysis from wildtype (WT) mice that received either WT bone marrow (n=9) or sickle cell disease bone marrow (n=7). c, MEL differentiation with 2% DMSO for 4 days shows increased *β-globin* expression assessed by qPCR relative to housekeeping gene *β-actin,* reflective of increased hemoglobin production. d, K562 differentiation with 1.5mM sodium butyrate for 4 days shows increased *β-GLOBIN* expression assessed by qPCR relative to housekeeping gene *β-ACTIN*, reflective of increased hemoglobin production. The statistical significance test for all graphs was done with an unpaired two-tailed t-test with a significance threshold level of 0.05.

**Extended Data Fig. 2:**
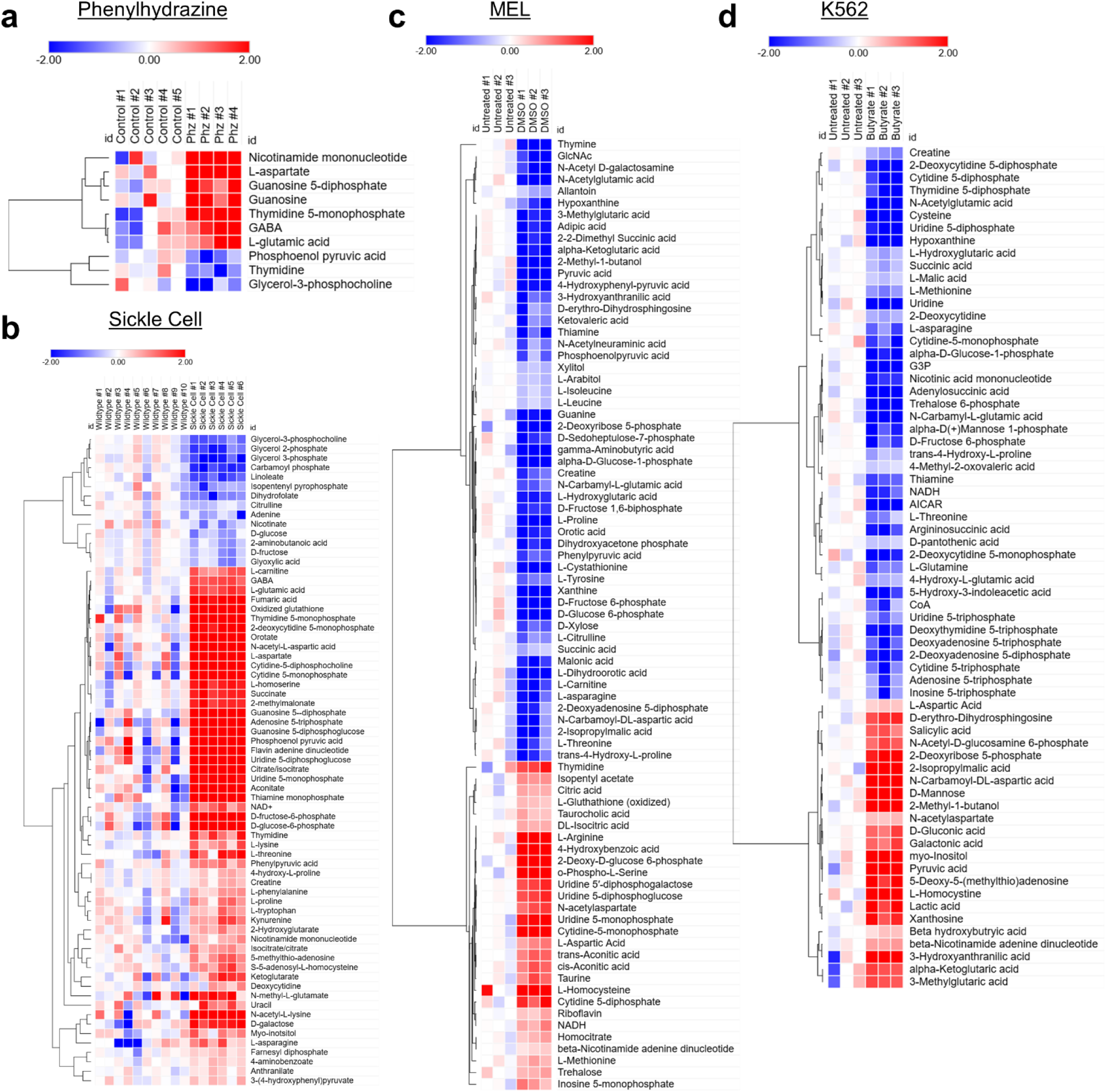
Metabolomic analysis of the four models of erythropoiesis. Heat map of the log_2_ of the median centered metabolite abundances in the (**a**) circulating RBCs of control mice and mice treated with phenylhydrazine (Phz), (**b**) circulating RBCs from wildtype mice that received either wildtype bone marrow or sickle cell disease bone marrow, (**c**) untreated and DMSO treated cell pellets of MEL cells, and (**d**) untreated and sodium butyrate treated cell pellets of K562 cells. Only significantly changed metabolites are shown (p-value < 0.05).

**Extended Data Fig. 3.**
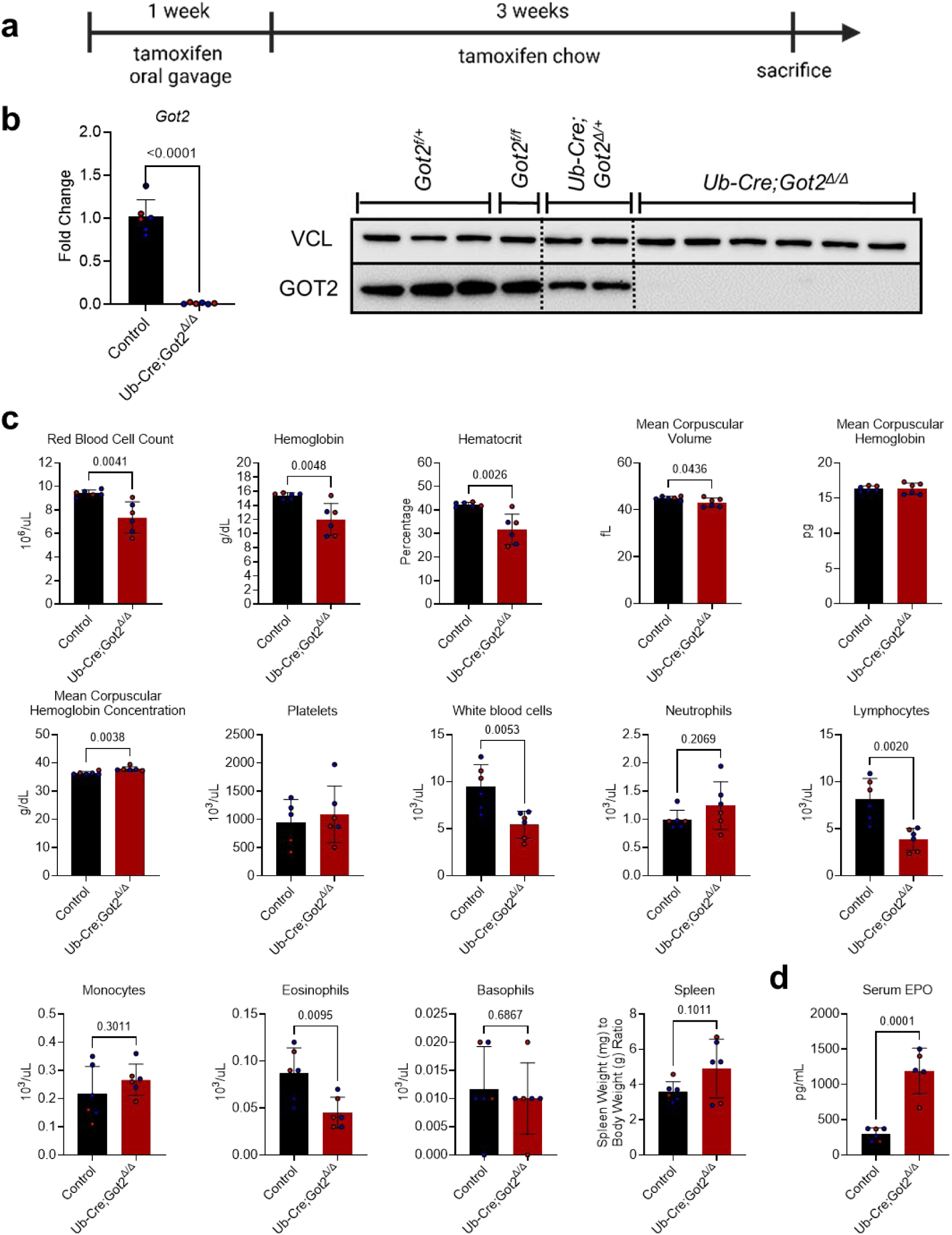
Ubiquitous, whole-body deletion of *Got2* in adult mice results in anemia. a, Experimental timeline of the control and *Ub-Cre;Got2^Δ/Δ^* mice experiment where all 8-12 week old mice were started on tamoxifen. **b,** *Got2* mRNA and GOT2 protein levels were analyzed in liver cells, demonstrating complete or near-complete absence of *Got2* expression in *Ub-Cre;Got2^Δ/Δ^* mice. *Got2* heterozygosity does not cause an erythroid defect, so control mice comprise the following genotypes: *Got2^f/+^, Got2^f/f^,* and *Ub-Cre;Got2^Δ/+^.* **c,** The complete blood counts and spleen weight of control (n=6) and *Ub-CreERT2;Got2^Δ/Δ^*mice (n=6) after 1 month of tamoxifen treatment. **d,** Elevated serum EPO levels and spleen size indicate appropriate EPO and extramedullary response to anemia in *Ub-CreERT2;Got2^Δ/Δ^* mice. Red and blue circles represent female and male mice, respectively. The statistical significance test for all graphs was done with an unpaired two-tailed t-test with a significance threshold level of 0.05.

**Extended Data Fig. 4:**
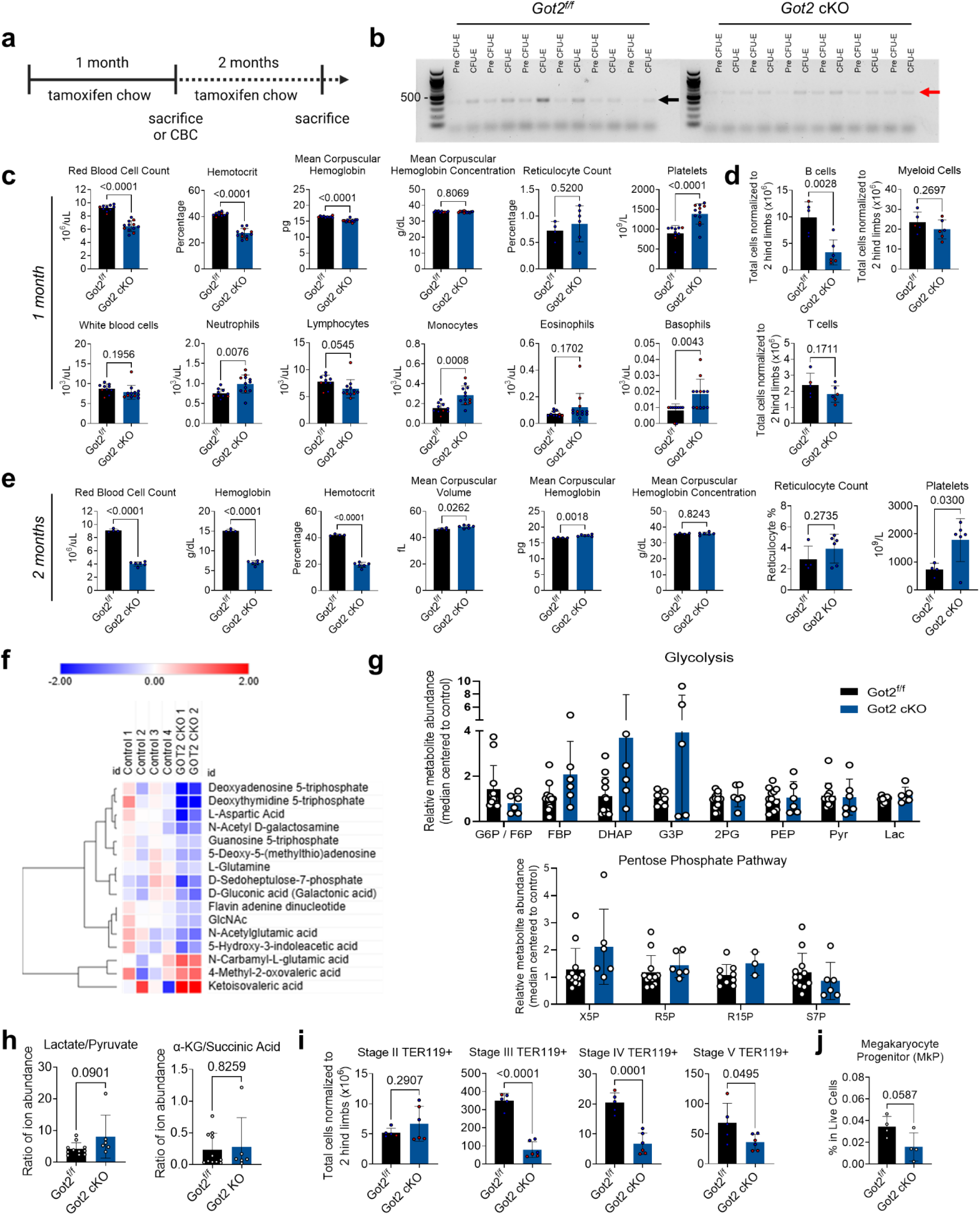
Blood and bone marrow analysis of erythroid-specific *Got2* cKO mice. a, Experimental timeline of the control and *Gata1-Cre;Got2^Δ/Δ^* (*Got2* cKO) mice experiment where all 5-8 week old mice were started on tamoxifen. Mice were either (i) sacrificed at 1 month for analysis or (ii) had a CBC done at 1 month and sacrificed at 2 months for analysis. **b,** Pre CFU-E or CFU-E cells from each mouse were flow sorted with the gating scheme shown in **Extended Data** Fig. 5, and DNA was isolated for PCR genotyping using primers listed in Supplementary Table 2. The floxed *Got2* allele is represented by a 385 base pair product (depicted by black arrow) and the *Got2* KO allele is represented by a 601 bp product (depicted by red arrow). **c,** The remaining red blood cell, white blood cell, and myeloid parameters in the blood of control (n=6) and *Got2* cKO mice (n=6) after 1 month of tamoxifen. Data presented are from 2 independent experiments. **d,** Total counts of B cells (B220 and CD19 positive), T cells (CD3e positive), and myeloid cells (CD11b and Gr1 positive) in the bone marrow of control and *Got2* KO mice after 1 month of tamoxifen. **e,** Complete blood counts control (n=4) and *Got2* cKO mice (n=5) after 2 months of tamoxifen. **f,** Heat map of the log_2_ of the median centered metabolite abundances (only significantly changed metabolites are shown (p-value < 0.05). **g,** Relative metabolite abundance in the indicated glycolytic and pentose phosphate pathway metabolites. Data presented are from 3 independent experiments. G6P: glucose-6-phosphate; F6P: fructose-6-phosphate; FBP: fructose-1,6-bisphosphate; DHAP: dihydroxyacetone phosphate; GA3P: glyceraldehyde-3-phosphate; 2PG: 2-phosphoglycerate; PEP: phosphoenol pyruvate; Pyr: pyruvate; Lac: Lactate; X5P: xylulose-5-phosphate; R5P: ribose-5; R15P: ribose-1,5-bisphosphate; S7P: seduoheptulose-7-phosphate. **h,** Relative ratios of the indicated metabolites in control and *Got1* cKO mice. α-KG: α-ketoglutarate. Data presented are from 3 independent experiments. **i,** Total counts of stage II (BasoEbs), III (PolyEbs), IV (OrthoEbs and immature reticulocytes), or V (mature RBCs) TER119+ cells (terminal erythroblasts) in the bone marrow. **j,** Frequency of the MkP population, which gives rise to platelets. Red and blue circles represent female and male mice, respectively. The statistical significance test for all graphs was done with an unpaired two-tailed t-test with a significance threshold level of 0.05.

**Extended Data Fig. 5:**
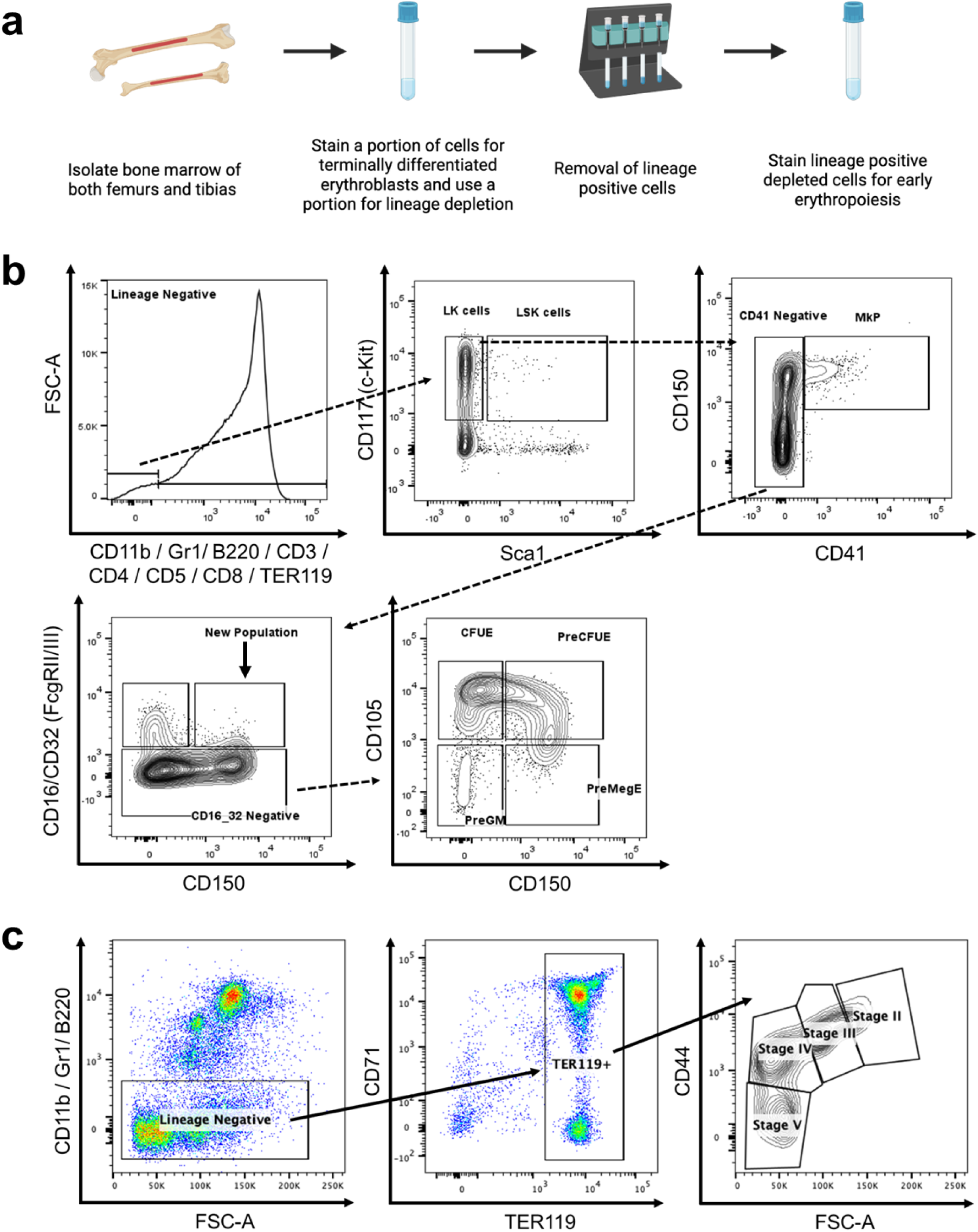
Flow gating strategy of bone marrow for cells of early erythropoiesis and terminal erythroid differentiation. a, Workflow for bone marrow isolation of 2 femurs and 2 tibias for flow analysis. b, Gating scheme for early erythropoiesis: Pre CFU-E, which encompass BFU-E and early CFU-E cells, are Lin^-^c-Kit^+^Sca1^-^CD41^-^CD16/CD32^-^CD105^+^CD150^+^. CFU-E cells are Lin^-^c-Kit^+^Sca1^-^CD41^-^CD16/CD32^-^ CD105^-^CD150^-^. The megakaryocyte progenitor (MkP) cells can also be isolated by gating on Lin^-^c-Kit^+^Sca1^-^ CD41^-^CD150^+^CD41^+^. c, Gating scheme for terminal erythroblasts: stage II are representative of basophilic erythroblasts, stage III are polychromatic erythroblasts, stage IV are orthochromatic erythroblasts with immature reticulocytes, and stage V are mostly mature RBCs.

**Extended Data Fig. 6:**
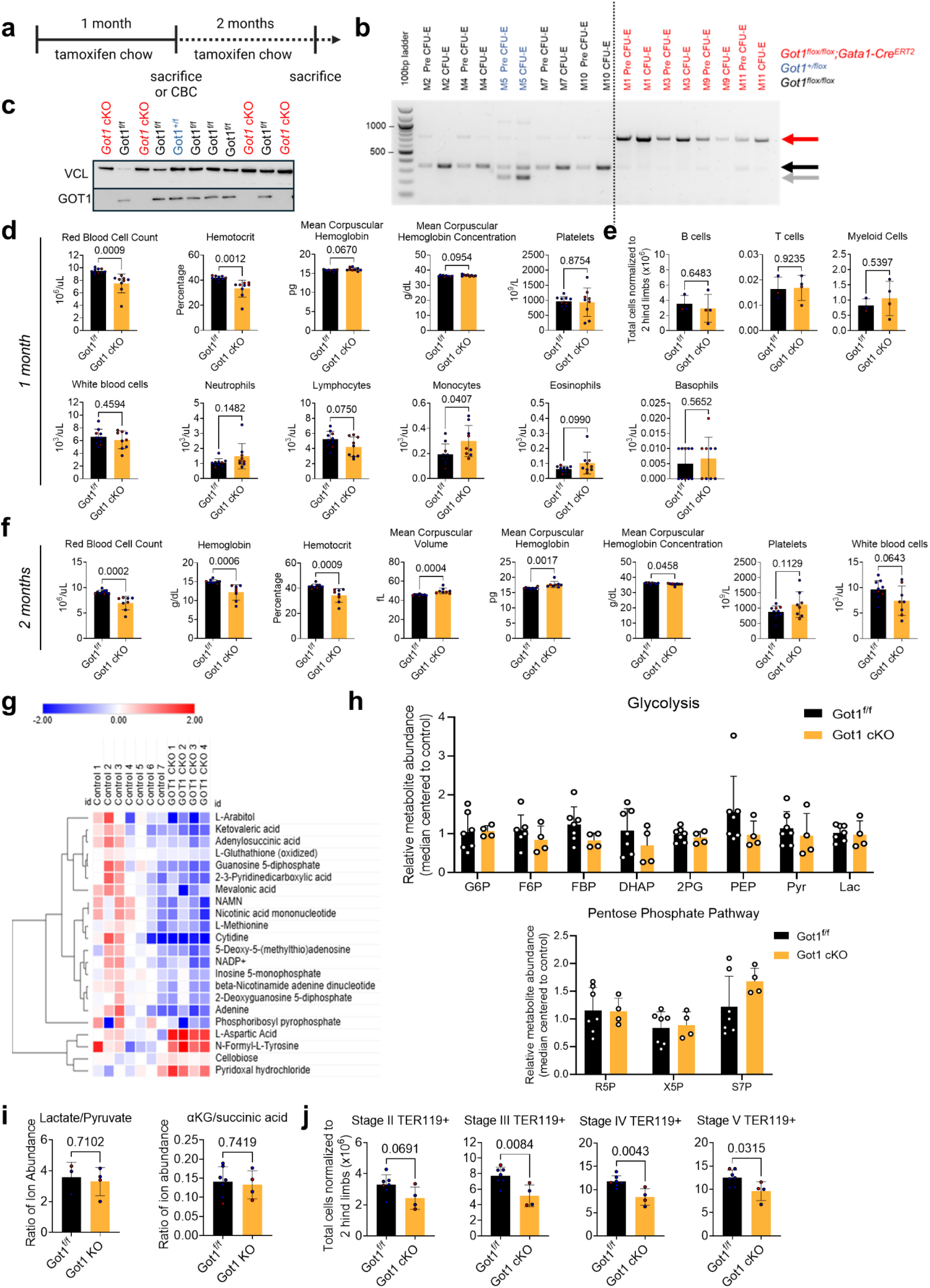
Blood and bone marrow analysis of erythroid-specific *Got1* cKO mice. a, Timeline of the control and *Gata1-Cre;Got1^Δ/Δ^* (*Got1* cKO) mice experiment, where all 6-10 week old mice were started on tamoxifen. Mice were either (i) sacrificed at 1 month for analysis or (ii) had a CBC done at 1 month and sacrificed at 2 months for analysis. **b,** Pre CFU-E or CFU-E cells from each mouse were flow sorted with the gating scheme shown in **Extended Data** Fig. 5B and DNA was isolated for PCR genotyping using primers listed in **Supplementary Table 2**. The wildtype allele is represented by a 250 base pair product (depicted by gray arrow), floxed *Got1* allele is represented by a 347 base pair product (depicted by black arrow) and the *Got1* KO allele is represented by a 749 bp product (depicted by red arrow). **c,** Western blot of GOT1 and VINCULIN loading control in circulating RBCs of control (Got1^+/f^, depicted in blue font, and Got1^f/f^, depicted in black font) and *Gata1-Cre;Got1^Δ/Δ^* mice (*Got1* KO, depicted in red font), showing that GOT1 is depleted. **d,** The remaining red blood cell, white blood cell, and myeloid parameters in the blood of the control (n=10) and *Got1* cKO mice (n=8) after 1 month of tamoxifen. Data presented are from 2 independent experiments. **e,** Total counts of B cells (B220 and CD19 positive), T cells (CD3e positive), and myeloid cells (CD11b and Gr1 positive) in the bone marrow of control and *Got1* cKO mice after 1 month of tamoxifen. **f,** Complete blood counts of control (n=10) and *Got1* cKO mice (n=8) after 2 months of tamoxifen. Data presented are from 2 independent experiments. **g,** Heat map of the log_2_ of the median centered metabolite abundances. Only significantly changed metabolites are shown (p-value < 0.05). **h,** Relative metabolite abundance in the indicated glycolytic and pentose phosphate pathway metabolites. G6P: glucose-6-phosphate; F6P: fructose-6-phosphate; FBP: fructose-1,6-bisphosphate; DHAP: dihydroxyacetone phosphate; 2PG: 2-phosphoglycerate; PEP: phosphoenol pyruvate; Pyr: pyruvate; Lac: Lactate; X5P: xylulose-5-phosphate; R5P: ribose-5-phosphate; S7P: sedoheptulose-7-phosphate. **i,** Relative ratios of the indicated metabolites in control and *Got1* cKO mice. α-KG: α-ketoglutarate. **j,** Total counts of stage II (BasoEbs), III (PolyEbs), IV (OrthoEbs and immature reticulocytes), or V (mature RBCs) TER119+ cells (terminal erythroblasts) in the bone marrow. Red and blue circles represent female and male mice, respectively. The statistical significance test for all graphs was done with an unpaired two-tailed t-test with a significance threshold level of 0.05.

**Extended Data Fig. 7:**
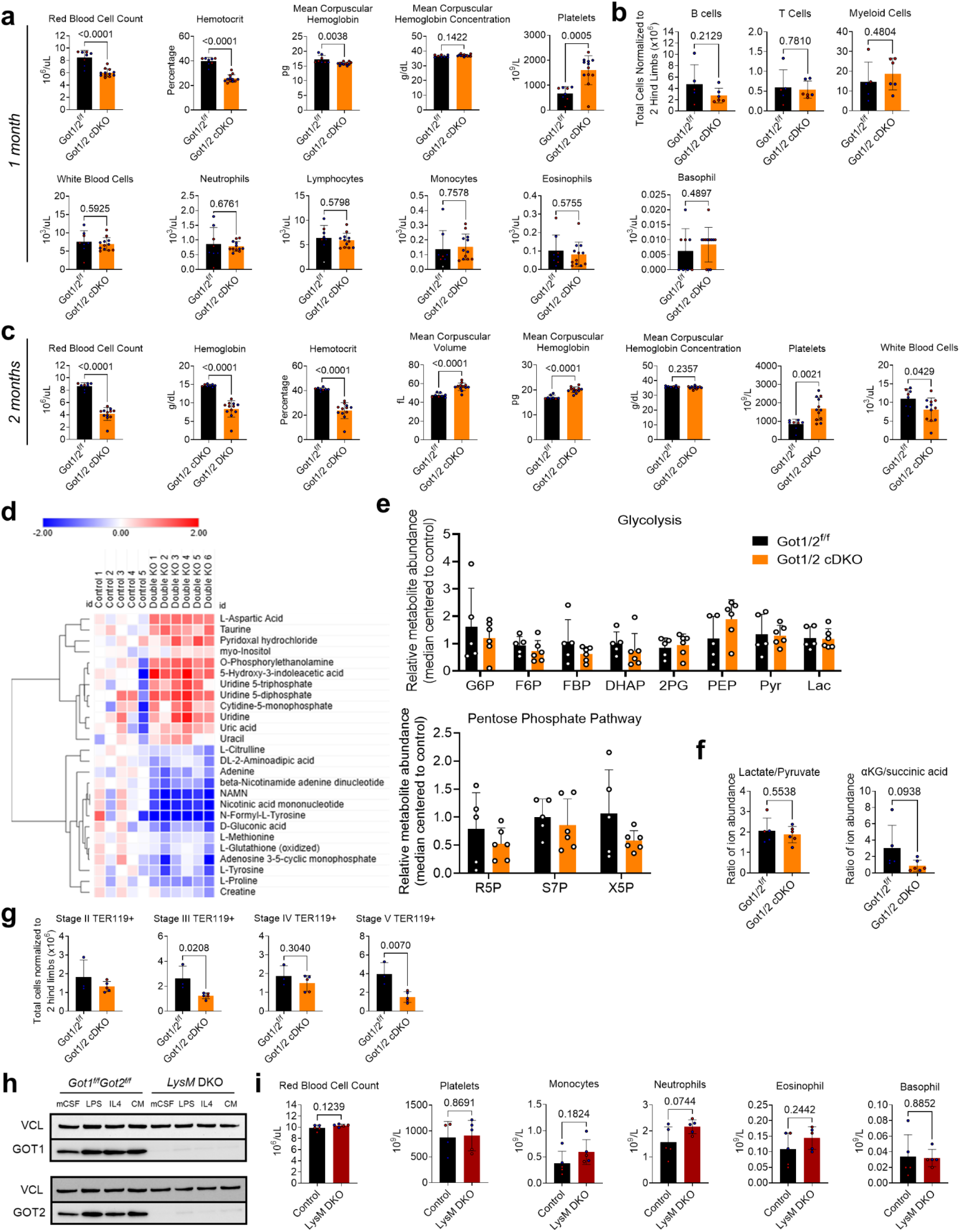
Blood and bone marrow analysis of erythroid-specific double *Got1/2* cKO mice and blood analysis of myeloid-specific double *Got1/2* cKO mice. a, The remaining red blood cell, white blood cell, and myeloid parameters in the blood of the 5-10 week old control (n=8) and *Gata1-CreERT2;Got1^Δ/Δ^Got2^Δ/Δ^*(*Got1/2* cDKO) mice (n=12); all mice received 1 month of tamoxifen. Data presented are from 2 independent experiments. b, Total counts of B cells (B220 and CD19 positive), T cells (CD3e positive), and myeloid cells (CD11b and Gr1 positive) in the bone marrow of control and *Got1/2* cDKO mice after 1 month of tamoxifen. c, Complete blood counts after 2 months of tamoxifen. Data presented are from 2 independent experiments. d, Heat map of the log_2_ of the median centered metabolite abundances. Only significantly changed metabolites are shown, p-value < 0.05. e, Relative metabolite abundance in the indicated glycolytic and pentose phosphate pathway metabolites. G6P: glucose-6-phosphate; F6P: fructose-6-phosphate; FBP: fructose-1,6-bisphosphate; DHAP: dihydroxyacetone phosphate; 2PG: 2-phosphoglycerate; PEP: phosphoenolpyruvate; Pyr: pyruvate; Lac: Lactate; X5P: xylulose-5-phosphate; R5P: ribose-5-phosphate; S7P: sedoheptulose-7-phosphate. f, Relative ratios of the indicated metabolites in control and *Got1/2* cDKO mice. α-KG: α-ketoglutarate. g, Total counts of stage II (BasoEbs), III (PolyEbs), IV (OrthoEbs and immature reticulocytes), or V (mature RBCs) TER119+ cells (terminal erythroblasts) in the bone marrow. h, Western blot of GOT1, GOT2 and VINCULIN loading control in isolated macrophages from the bone marrow of *Got1^f/f^Got2^f/f^*control mice or *LysM-Cre;Got1^Δ/Δ^Got2^Δ/Δ^* (*LysM* cDKO) mice that were then cultured in M-CSF, LPS, IL-4, or condition media (CM) from tumors, showing preserved absence of protein within various macrophage states. i, Red blood cell, monocytes and granulocytes (neutrophils, eosinophils, and basophils) of control (*Got1^f/f^Got2^f/f^,* n=5) and *LysM-Cre;Got1^Δ/Δ^Got2^Δ/Δ^*(*LysM* cDKO, n=5) mice. Red and blue circles represent female and male mice, respectively. The statistical significance test for all graphs was done with an unpaired two-tailed t-test with a significance threshold level of 0.05.

**Extended Data Fig. 8:**
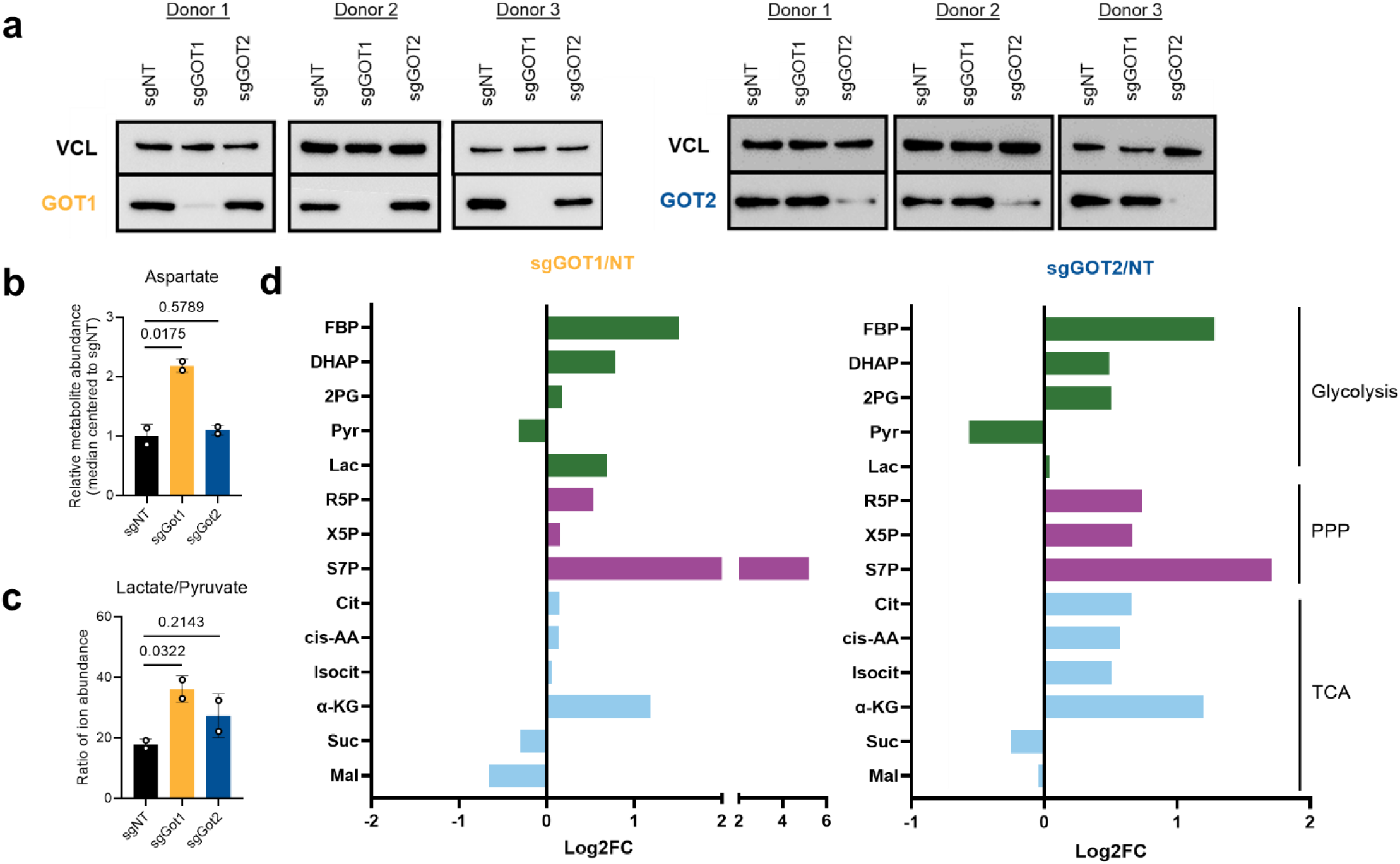
*GOT* knockout leads to NADH reductive stress in human erythroid progenitor cells. a, Western blot of GOT1, GOT2 and VINCULIN loading control in HSPC-derived erythroid cells at day 11 of differentiation. Data from 3 different human donors are shown. **b,** Relative abundance of aspartate in day 14 erythroid cells. **c,** Relative ratio of lactate/pyruvate in HSPC-derived erythroid cells at day 14 of differentiation. **d,** Relative fold changes in the indicated glycolytic, pentose phosphate pathway and tricyclic acid cycle metabolites at day 14 of differentiation. G6P: glucose-6-phosphate; F6P: fructose-6-phosphate; FBP: fructose-1,6-bisphosphate; DHAP: dihydroxyacetone phosphate; G3P: glyceraldehyde-3-phosphate; PEP: phosphoenol pyruvate; Pyr: pyruvate; Lac: Lactate; X5P: xylulose-5-phosphate; R5P: ribose-5-phosphate; R15P: ribose-1,5-bisphosphate; S7P: seduoheptulose-7-phosphate; Cit: citrate; cis-AA: cis-aconitate; Isocit: isocitrate; α-KG: α-ketoglutarate; Suc: succinate; Mal: malate.

**Extended Data Fig. 9:**
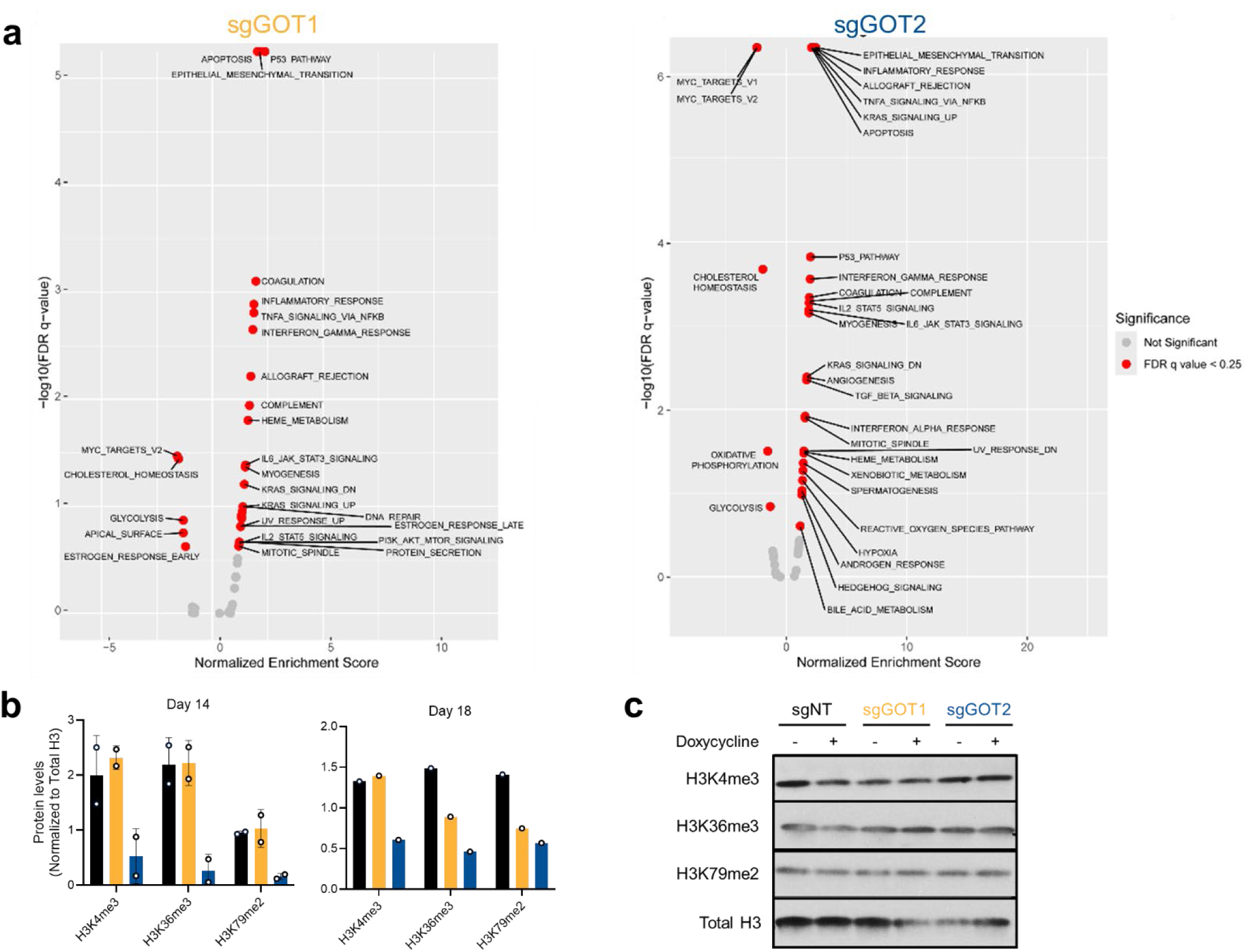
*GOT* knockout leads to increased apoptotic pathways in human erythroid progenitor cells. a, Top enriched signatures associated with up-regulated or down-regulated genes in *GOT1* (left) or *GOT2* (right) KO HSPC-derived erythroid cells at day 5 of differentiation. The data is derived using three human donors. FDR, false discovery rate. **b,** Densitometry quantification of histone bands in the Western blot shown in Fig. 4F. **c,** Western blot of H3K4me3, H3K36me3, H3K79me2 and total H3 loading control in MIAPaCa-2 human pancreatic cancer cell lines with doxycycline-inducible nontargeting (NT), Got1 or Got2 shRNA.

**Supplemental Table 1.**
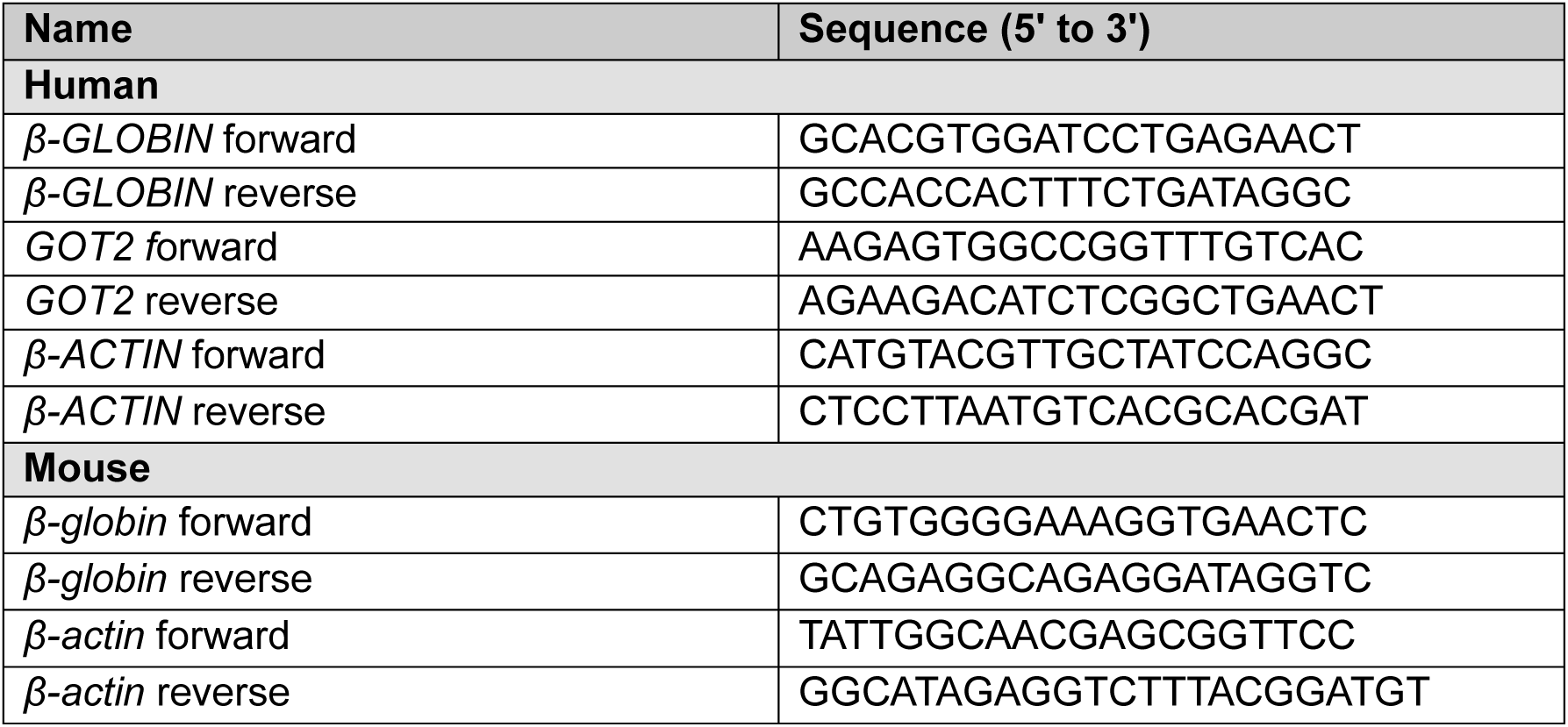
List of qPCR primers.

**Supplemental Table 2.**
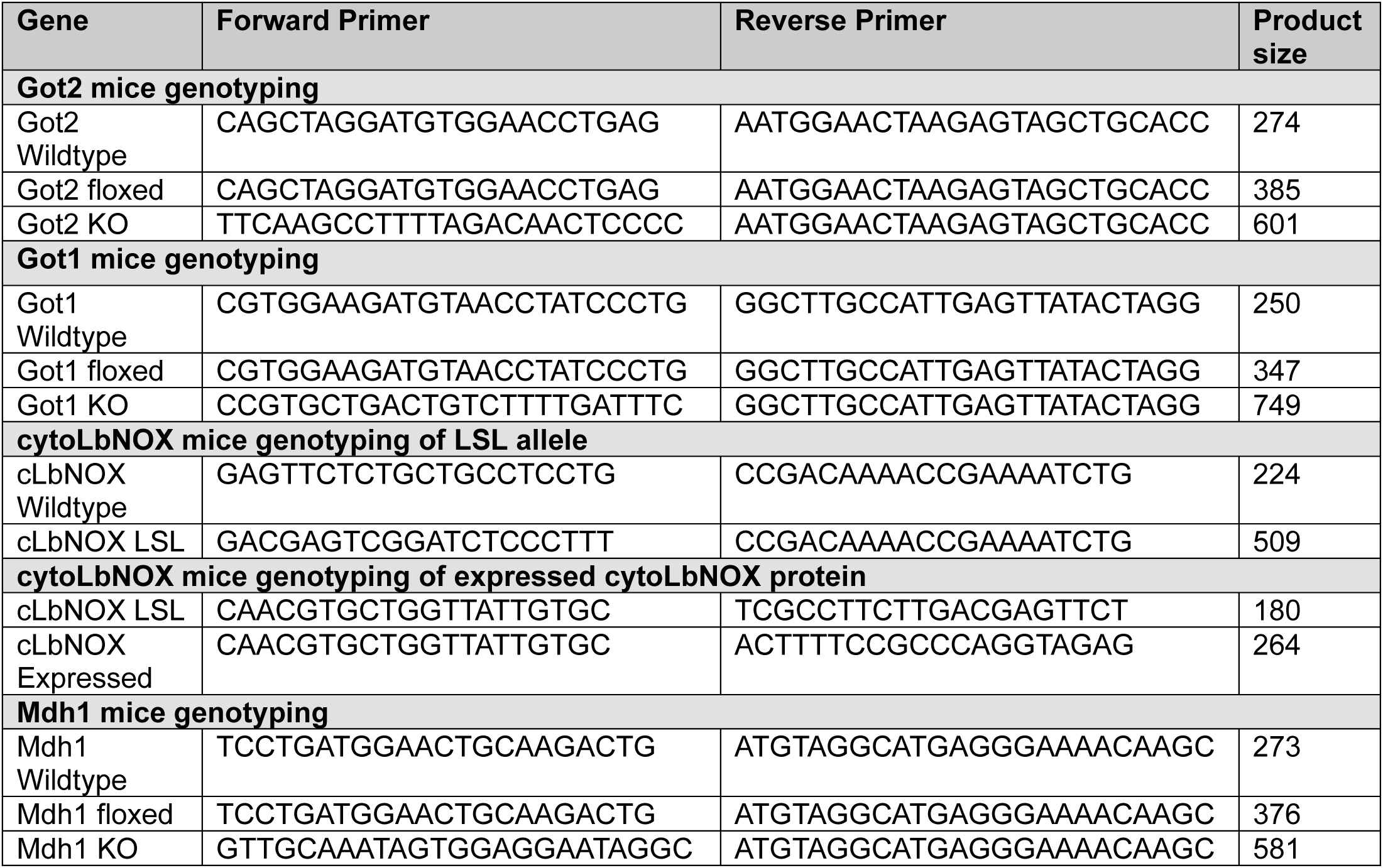
List of PCR genotyping primers.

**Supplemental Table 3.** List of antibodies and the reagent suppliers.

| Human (Flow) | Fluorescence | Catalog Number | Vendor |
| --- | --- | --- | --- |
| CD235a (Glycophorin A) | APC | 551336 | BD Biosciences |
| CD71 | PE | 334106 | BioLegend |
| CD233 (Band 3) | FITC | 9439FI | IBGRL |
| CD49d ($\alpha_4$-integrin) | PE-Cy7 | 304314 | Biolegend |
| CD11b | PE | 301305 | BioLegend |
| Mouse (Flow) | Fluorescence | Catalog Number | Vendor |
| **Early Erythropoiesis** |  |  |  |
| Sca1 | BV421 | 108128 | Biolegend |
| c-Kit / CD117 | APC-Cy7 | 105825 | BioLegend |
| CD41 | Alexa Fluor 700 | 133925 | BioLegend |
| CD150 | APC | 115910 | BioLegend |
| CD16/32 | BV605 | 563006 | BD Biosciences |
| CD105 / Endoglin | PE-Cy7 | 120410 | BioLegend |
| CD11b | Biotin | 101208 | Biolegend |
| Gr1 | Biotin | 108408 | Biolegend |
| B220 | Biotin | 103208 | Biolegend |
| CD3 | Biotin | 100308 | Biolegend |
| CD4 | Biotin | 116006 | Biolegend |
| CD5 | Biotin | 100608 | Biolegend |
| CD8 | Biotin | 100708 | Biolegend |
| TER119 | Biotin | 116208 | Biolegend |
| PE-Cy5 | Streptavidin |  |  |
| **Terminal erythroid differentiation** |  |  |  |
| CD71 | PE-Dazzle | 113818 | BioLegend |
| TER119 | PE-Cy7 | 116222 | BioLegend |
| CD44 | APC | 103012 | BioLegend |
| B220 | APC-Cy7 | 103224 | BioLegend |
| Gr1 | APC-Cy7 | 108424 | Biolegend |
| CD11b | APC-Cy7 | 101226 | Biolegend |
| **B/T/Myeloid** |  |  |  |
| Gr1 | PE | 108408 | Biolegend |
| CD19 | PE-Cy7 | 115520 | Biolegend |
| B220 | FITC | 103206 | Biolegend |
| CD3e | APC | 17-0031-82 | Invitrogen |
| Live/Dead (Flow) | Fluorescence | Catalog Number | Vendor |
| DAPI | Pacific Blue | 422801 | BioLegend |
| Zombie Aqua | BV510 | 423101 | Biolegend |
| Annexin V | AF488 | A13201 | Invitrogen |
| Annexin V | Pacific Blue | A35122 | Invitrogen |
| Human and Mouse (Western) | Dilution | Catalog Number | Vendor |
| Vinculin | 1:10,000 | 13901 | Cell Signaling |
| GOT1 | 1:1,000 | ab171939 | Abcam |
| GOT2 | 1:1,000 | HPA018139 | Sigma |
| MDH1 | 1:6,000 | ab180152 | Abcam |
| H3K4me3 | 1:1,000 | 9751 | Cell Signaling |
| H3K36me3 | 1:1,000 | 61021 | Activ Motif |
| H3K79me2 | 1:1,000 | 5427 | Cell Signaling |
| Total H3 | 1:5,000 | 3638 | Cell Signaling |

